# Impaired redox and protein homeostasis as risk factors and therapeutic targets in toxin-induced biliary atresia

**DOI:** 10.1101/821967

**Authors:** Xiao Zhao, Kristin Lorent, Diana Escobar-Zarate, Ramakrishnan Rajagopalan, Kathleen M. Loomes, Kevin Gillespie, Clementina Mesaros, Michelle A. Estrada, Ian Blair, Jeffrey D. Winkler, Nancy B. Spinner, Marcella Devoto, Michael Pack

## Abstract

**BACKGROUND and AIMS:** Extra-hepatic biliary atresia (BA) is a pediatric liver disease with no approved medical therapy. Recent studies using human samples and experimental modeling suggest that glutathione redox metabolism and heterogeneity play a role in disease pathogenesis. We sought to dissect the mechanistic basis of liver redox variation and explore how other stress responses affect cholangiocyte injury in BA.

**METHODS:** We performed quantitative *in situ* hepatic glutathione redox mapping in zebrafish larvae carrying targeted mutations in glutathione metabolism genes and correlated these findings with sensitivity to the plant-derived BA-linked toxin biliatresone. We also determined whether genetic disruption of HSP90 protein quality control pathway genes implicated in human BA altered biliatresone toxicity in zebrafish and human cholangiocytes. An *in vivo* screen of a known drug library was performed to identify novel modifiers of cholangiocyte injury in the zebrafish experimental BA model with subsequent validation.

**RESULTS:** Glutathione metabolism gene mutations caused regionally distinct changes in the redox potential of cholangiocytes that differentially sensitized them to biliatresone. Disruption of human BA-implicated HSP90 pathway genes sensitized zebrafish and human cholangiocytes to biliatresone-induced injury independent of glutathione. Phosphodiesterase-5 inhibitors (PDE5i) and other cGMP signaling activators worked synergistically with the glutathione precursor *N-* acetylcysteine (NAC) in preventing biliatresone-induced injury in zebrafish and human cholangiocytes. PDE5i enhanced proteasomal degradation and required intact HSP90 chaperone.

**CONCLUSION:** Regional variation in glutathione metabolism underlies sensitivity to the biliary toxin biliatresone, and mirrors recently reported BA risk stratification linked to glutathione metabolism gene expression. Human BA can be causatively linked to genetic modulation of protein quality control. Combined treatment with NAC and cGMP signaling enhancers warrants further investigation as therapy for BA.

**What You Need to Know:** *Background and Context:* Biliary atresia (BA) is an obstructive fibrosing cholangiopathy that is the leading indication for liver transplantation in the pediatric population. There are no known treatments to prevent progressive liver injury after surgical restoration of bile flow.

*New Findings:* The authors identify factors that affect susceptibility of cholangiocytes to oxidative injury using a toxin-induced BA model. This information is used to validate genetic risk factors for human BA and identified PDE5i as a potential treatment for biliary atresia, either on its own or in combination with the anti-oxidant *N-*acetyl-cysteine.

*Limitations:* The work done in animal and cell culture models needs further study in human tissue-derived models and a larger cohort of BA patients.

*Impact:* The findings from this study provide a rationale for identifying new genetic risk factors that predispose to BA and for an interventional study to prevent progressive liver injury in this enigmatic disease.

*Short Summary:* This study uses zebrafish and human cell culture models to identify novel injury mechanisms, genetic risk factors and new therapies for the pediatric liver disease biliary atresia.

---

Extra-hepatic biliary atresia (BA) is a fibrosing cholangiopathy of infancy that is the most common indication for pediatric liver transplantation.^1, 2^ Although genetic etiology has been implicated in syndromic BA,^3^ genetic factors have not been conclusively identified for the majority of cases, which occur in the absence of other defects (referred to as isolated BA). Indeed, the most commonly accepted mechanistic model posits that isolated BA results from late gestational exposure of genetically susceptible individuals to an environmental trigger, such as a toxin or virus,^4,5^ thus highlighting the importance of studying BA in the context of genetic-environment interactions.

Although suspected, environmental triggers for human BA have yet to be definitively identified. Naturally-occurring BA epidemics in newborn livestock, however, have been linked to maternal ingestion of biliatresone, a highly reactive isoflavone electrophile identified from *Dysphania* plant species that forms adducts with reduced glutathione (GSH), nucleic acids and amino acids *in vitro*.^6,7^ Despite its reactivity, *in vivo* biliatresone exposure is selectively toxic toward extrahepatic cholangiocytes (EHC), sparing intrahepatic cholangiocytes (IHC) and hepatocytes. Prior work in the zebrafish model causatively linked EHC susceptibility to their low redox reserve compared to the resistant cells. Consistent with this model, treatment of zebrafish larvae with *N-*acetylcysteine (NAC), a glutathione precursor used for treatment of acetaminophen overdose,^8^ temporarily delayed the onset of biliatresone-induced EHC injury, whereas pharmacological depletion of glutathione sensitized IHC to the toxin.^9^

Although direct consumption of biliatresone by humans is unlikely, our findings from the zebrafish model were the first to argue that variations in redox metabolism could be a factor driving human BA pathogenesis. Supporting this idea, high hepatic expression of glutathione metabolism genes was recently shown to be positively associated with transplant-free survival in infants with BA following Kasai portoenterostomy. NAC treatment also reduced liver injury and fibrosis in the rhesus rotavirus murine BA model.^10^

To gain a better understanding of the factors affecting regional variations in cholangiocyte redox metabolism and how it might influence BA susceptibility and outcomes, we took advantage of the genetic and drug-screening capabilities of the zebrafish system to identify modifiers of biliatresone toxicity. These studies showed that EHC and IHC depend on distinct arms of the glutathione metabolism pathway to maintain redox homeostasis. Our work further revealed that protein quality control (PQC) mechanisms work in concert with redox responses following biliatresone exposure, and genetic disruption of the heat shock chaperone 90 (HSP90) pathway genes (*STIP1, REV1*), implicated in human BA *via* whole exome sequencing,^11^ altered biliatresone toxicity in the zebrafish and human cholangiocytes. Finally, we identified pharmacological activators of cGMP signaling as a class of drugs that work synergistically with NAC in attenuating biliatresone-mediated injury by enhancing PQC. These data highlight the complex nature of BA pathogenesis through demonstrating roles for both redox and protein homeostasis in cholangiocyte injury responses, and identified a novel treatment strategy that target both pathways.

## Materials and Methods

### Fish care and strains

The wild-type (WT) strains (TL, AB), transgenic strains (Tg (*ef1a*:Grx-roGFP; Tg(*Tp1*:mCherry), (*Tg(fabp10a:TRAP)*; (*Tg(krt18:TRAP)*) and mutant strains (*gclm, abcc2, gsr, stip1, rev1*) were used for the present studies. All zebrafish experiments were approved by the Institutional Animal Care and Use Committee at the University of Pennsylvania Perelman School of Medicine.

### Generation of targeted mutations in zebrafish using CRISPR/Cas9 system

The target sequences for *gclm, abcc2, stip1, rev1* were identified using the website: https://chopchop.cbu.uib.no. The single-guide RNA was generated as described.^12^ Cas9 protein was obtained from PNABIO. Approximately 60 to 120 pg of sgRNA and 180 pg of Cas9 protein were co-injected into WT embryos at the one-cell stage. Details on subsequent genotyping are described in the supplementary methods. All experiments were performed on the animals of F3 or subsequent generations. The genotyping primers for *gclm, abcc2, stip1, rev1* mutants are listed in Table S1. The *gsr* mutant fish were genotyped using the KASP genotyping assays (KBioscience).

### Redox-Sensitive GFP Redox Mapping

Double transgenic (*Tg(ef1a:Grx-roGFP; tp1:mCherry*)) larvae in *gclm, abcc2, gsr* genetic background were used for the redox mapping of hepatocytes, IHCs, and EHCs. Conservation of the biosensor redox state was achieved with the thiol-alkylating agent, *N-*ethylmaleimide (NEM), and tissue fluorescence redox imaging was performed as previously described.^9, 13^

### GSH Derivatization and Liquid Chromatography/Mass Spectrometry

The GSH content was measured in larval livers of WT and mutant fish treated with biliatresone *vs* DMSO as previously described.^9^ Five larval livers (including the extrahepatic biliary system) were used per sample.

### Cell culture

H69 cells, an SV40-immortalized normal human cholangiocyte cell line, and low passage number normal human cholangiocytes (NHCs) were obtained from Nicholas F. LaRusso (Mayo Clinic) and grown in H69 media as previously described.^14^

### Drug Screen

5 dpf zebrafish larvae were arrayed in 100 μl E3 media in 96 well plates; 6 larvae per well containing individual compound from the Johns Hopkins University Clinical Compound Library (JHCC) (20μM) and biliatresone (0.5ug/ml) for 24 hr, followed by bodipy-C16 exposure for 4 hr. Biliatresone was synthesized as previously described.^15^ Fluorescent images of larval gallbladder within the 96-well plates were obtained using an Olympus inverted compound microscope *via* the Metamorph software. Compounds were considered active if gallbladder morphology appeared normal in 3 or more larvae per well. Compounds demonstrating toxicity were re-tested at either 10 μM or 5 μM final concentration. Each 96 well plate included two control wells with larvae that received no compound and two that were exposed to biliatresone alone.

### Drug Treatments

Zebrafish larvae (5 dpf) or cultured cholangiocytes were exposed to biliatresone and/or indicated compounds at the specified dose and duration. Details on the chemical treatments can be found in the supplementary methods. The medium containing each compound was changed daily. All assays were repeated three times.

***Immunofluorescence; Nanostring nCounter assay and data analysis; siRNA Transfection; RNA Extraction and qRT-PCR; Protein Extracts and Western Blots; Cell Viability Assay;***

### Proteasomal Degradation Assay

Details are described in the Supplementary Methods. Primer sequences are listed in Table S2.

### Statistical Analyses

Data represent at least three independent experiments reported as means +/- SEM. The Student t test was used for comparison between two groups. Data from three or more groups were analyzed by one-way analysis of variance (ANOVA) with Tukey’s multiple comparisons test. p < 0.05 was considered to be significant.

## Results

### Genetic disruption of glutathione metabolism genes in zebrafish

To gain insight into the differential sensitivity of zebrafish cholangiocytes to electrophilic stress, we analyzed the effect of biliatresone in larvae carrying loss-of-function mutations in glutathione metabolism genes. To study the role of glutathione synthesis and transport, we used CRISPR-Cas9 genome editing to disrupt *gclm*, which encodes the regulatory subunit of the rate-limiting enzyme in GSH synthesis, glutamate cysteine ligase (Gcl),^16^ and *abcc2*,^17^ which encodes the canalicular glutathione transporter, Mrp2. Mrp2 plays a central role in biliary GSH metabolism, as hepatocyte GSH secreted into bile is metabolized into its constituent amino acids and then resynthesized by cholangiocytes. Compared to the *gclm* mutation, which disrupts GSH synthesis in all cell types, mutation in *abcc2* specifically alters GSH synthesis in cholangiocytes. To study the role of GSH regeneration from its oxidized form GSSG,^18^ we analyzed larvae carrying a previously uncharacterized nonsense mutation in the *glutathione reductase* (*gsr*) gene (sa15125) that was generated *via* chemical mutagenesis (Fig. S1A, Table. S1).

Zebrafish homozygous for each mutation developed normally with normal fertility and lifespan (Fig. S1B-D’’’). The mutations were confirmed *via* genomic and cDNA sequencing, with quantitative RT-PCR showing significantly reduced levels of each targeted transcript, presumably as a result of nonsense-mediated mRNA decay (Fig. S1E). Mass spectrometry studies also confirmed that each mutation had the predicted effect on GSH levels, namely a pronounced reduction in the total hepatic GSH content of *gclm* mutants compared to their WT siblings, as well as a significant decrease in bile GSH measured in gallbladders dissected from adult *abcc2* mutants (Fig. S1F, G). Total hepatic GSH was not affected in *gsr* larvae, similar to *Gsr* mice (Fig. S1F).^19^

### Mutations of glutathione metabolism genes alter cellular redox status

To determine how mutations in glutathione metabolism genes affect the basal glutathione redox state of different liver cell types, we crossed the mutant lines into transgenic fish carrying a ubiquitously expressed cytoplasmic redox biosensor Grx-roGFP2 (hereafter, roGFP).^9, 13^ This highly sensitive and specific *in vivo* probe contains an engineered dithiol/disulfide switch that provides a ratiometric readout of fluorescence emission at two excitation wavelengths (405 and 488 nm). A high 405nm:488nm excitation ratio indicates an oxidized sensor (low GSH:GSSG ratio), whereas a low ratio signifies a reduced state (high GSH:GSSG ratio).

We found that homozygous *gclm* and *gsr* mutants exhibited an oxidized glutathione pool in all examined liver cell types (EHC, IHC and hepatocytes) compared to their wild-type (WT) and heterozygous sibling larvae, consistent with global disruption of GSH synthesis and regeneration, respectively (Fig. 1A-D). Predictably, the *abcc2* mutation led to oxidization of the glutathione pool in EHC and IHC, reflective of reduced GSH transport into bile, but had no effect on hepatocytes (Fig. 1E, F; S1G). Interestingly, the hepatic redox heterogeneity previously described^9^ was preserved in these mutant larvae with EHC remaining the most oxidized cell type at baseline compared to IHC and hepatocytes.

**Fig. 1.**
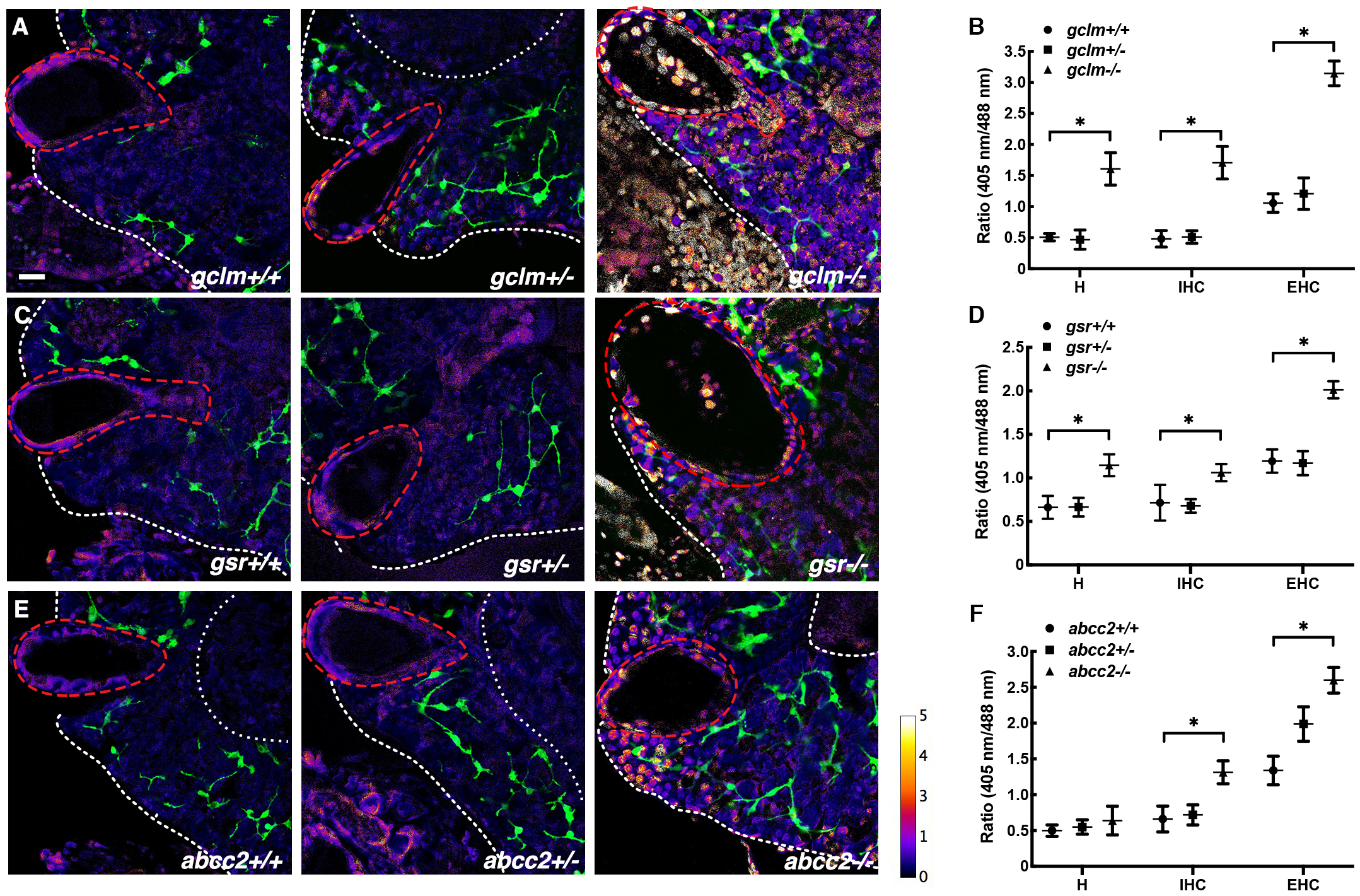
Glutathione metabolism gene mutations alter EHC, IHC and hepatocyte redox status. (A, C, E) *In situ* measurement of basal glutathione redox potential of 5 dpf *gclm* (A), *gsr* (C), *abcc2* (E) bigenic mutant larvae Tg (*e1f*: Grx1-roGFP; *tp1:*mCherry). Glutathione redox status of EHC (gallbladder-dashed red circle), IHC, and hepatocytes was derived from the roGFP 405nm/488nm fluorescence excitation ratio (higher excitation ratio-greater oxidation). EHC and hepatocytes localized by morphology. IHC localized by mCherry expression (pseudo-colored green). White dashed line - liver border. Scale bar, 10 μm. (B, D, F) Mean fluorescent 405 nm/488 nm excitation ratios in hepatocytes, IHC, and EHC for each genotype. n =10 larvae per genotype, means +/- SEM, *p<0.02 in comparison to WT hepatocytes, IHC, or EHC respectively.

### Changes in redox status reveal cell-type specific responses to electrophilic stress

Having defined the effects of glutathione metabolism gene mutations on basal redox status, we next asked whether they were predictive of sensitization to biliatresone in either EHC or IHC. We have previously shown that short duration exposure of larvae to biliatresone (4 hr) causes further oxidation of EHC, thus predicting overt injury that is first evident at 16-24 hr of continuous treatment.^9^ For this study, we used previously defined criteria for sensitization^9^ as severe EHC injury (complete to near complete gallbladder destruction) caused by a normally inactive low dose of the toxin (0.25 μg/ml, 24hr; Fig. 2A, A’, E), or IHC injury (hypoplastic intrahepatic ducts with short ductular projection) caused by either the low dose, or the standard dose (0.5 μg/ml, 24hr), which normally only induces EHC injury in WT larvae (Fig. 2A’’, F).

**Fig. 2.**
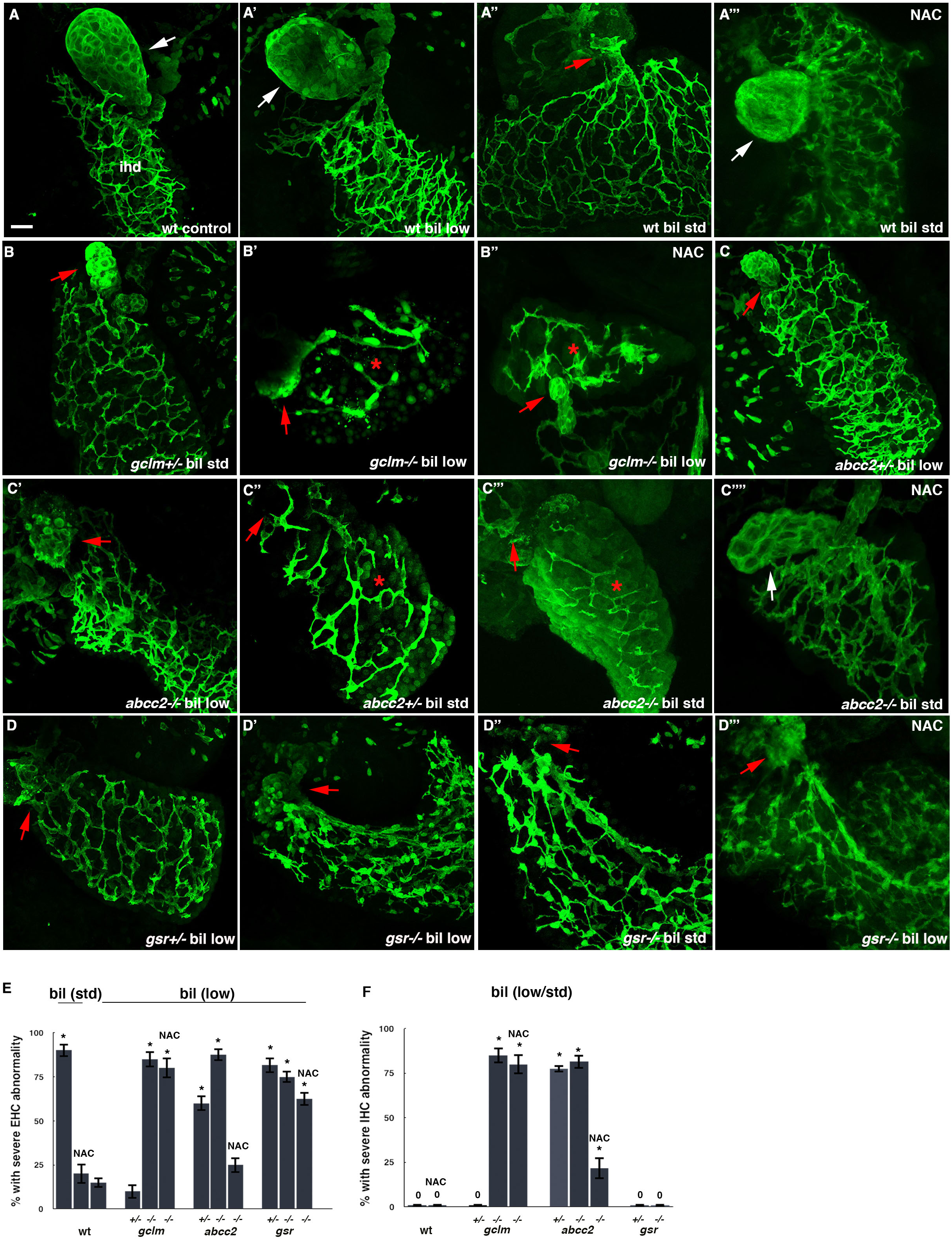
Glutathione metabolism gene mutations differentially sensitize cholangiocytes to biliatresone. (A-D’’’) 5 dpf WT (A-A’’’), *gclm* (B-B’’), *abcc2* (C-C’’’’), *gsr* (D-D’’’) mutant larvae were exposed to low-dose or standard dose biliatresone with or without NAC for 24 hr. Confocal projections through the livers of larvae immunostained with anti-Annexin A4 antibody show destruction of EHC and IHC at the specified genotype and dose exposure. White arrow – gallbladder; red arrow – gallbladder remnant. * denotes liver with IHC injury. Scale bar, 20 μm. (E & F) Percentage of larvae with severe EHC or IHC abnormalities at the indicated exposure(s) and genotype, respectively. *n* = 30-36 per genotype per condition, means +/- SEM, *p<0.01 in comparison to WT control EHC or IHC.

Previous studies of biliatresone in the zebrafish model suggested that oxidizing mutations would sensitize cholangiocytes to biliatresone.^9^ Consistent with these predictions, we observed that the oxidized EHC of *gclm, abcc2* and *gsr* mutants were all sensitized to low-dose biliatresone (Fig. 2B’, C, C’, D’, E). The oxidized IHC of *gclm* homozygotes and *abcc2* homozygotes were also sensitized to low and standard doses of the toxin, respectively (Fig. 2B’, C’’’, F). However, basal redox status was not always predictive of biliatresone sensitization. Specifically, we observed sensitization of IHC in *abcc2* heterozygotes and EHC in *gsr* heterozygotes, even though their basal redox potential was not significantly different from their WT siblings (Fig. 2C’’, D, E, F). Conversely, the IHC of *gsr* homozygotes were resistant (not sensitized) to biliatresone despite their oxidation (Fig. 2D’’, F). Collectively, these findings identify important differences in the way EHC and IHC maintain redox homeostasis in response to oxidative stress. EHC rely on GSH synthesis and regeneration, whereas GSH regeneration is not essential in IHC, presumably reflecting a greater synthetic capacity. Supporting this, the glutathione precursor NAC did not attenuate biliatresone-induced EHC injury in *gclm* or *gsr* mutants, but did rescue *abcc2* mutants. (Fig. 2A’’’, B’’, C’’’’, D’’’, E).

### Biliatresone induces proteomic stress and modulation of the Hsp90 chaperone machinery enhances biliatresone toxicity

Prior hepatic expression profiling experiments showed that in addition to glutathione metabolism genes, genes involved in the maintenance of cellular proteostasis were also upregulated in the initial response to biliatresone.^9^ Quantification of polysomal mRNA isolated from larval biliary cells (EHC & IHC) and hepatocytes *via* translating ribosomal affinity profiling (TRAP) methodology^19^ confirmed the induction of heat shock response (HSR) and unfolded protein response/endoplasmic reticulum (UPR/ER) stress response genes in both cell types 4 hr after biliatresone exposure (Fig. 3A). To test the functional significance of these findings, we asked whether an Hsp90 inhibitor 17-*N*-allylamino-17-demethoxygeldanamycin (17-AAG)^20^ sensitized WT larvae to low-dose biliatresone. Conversely, we also examined whether pre-treatment with an activator of Hsp70 (geranylgeranyl acetate),^21^ which functions upstream of Hsp90 chaperone function, could blunt biliatresone-mediated injury. We found that inhibition of Hsp90 sensitized EHC to low-dose biliatresone (Fig. 3B, B’, C, C’, E), whereas pre-treatment with the Hsp70 activator did not mitigate its toxic effect (Fig. 3B’’, D-E). Significantly, the GSH synthetic precursor NAC was unable to attenuate the sensitization caused by Hsp90 inhibition (Fig. 3C’’, E). Altogether, these data suggest that the Hsp70/Hsp90 chaperone machinery plays a crucial role in the cellular response to biliatresone that is independent of the glutathione-mediated redox response.

**Fig. 3.**
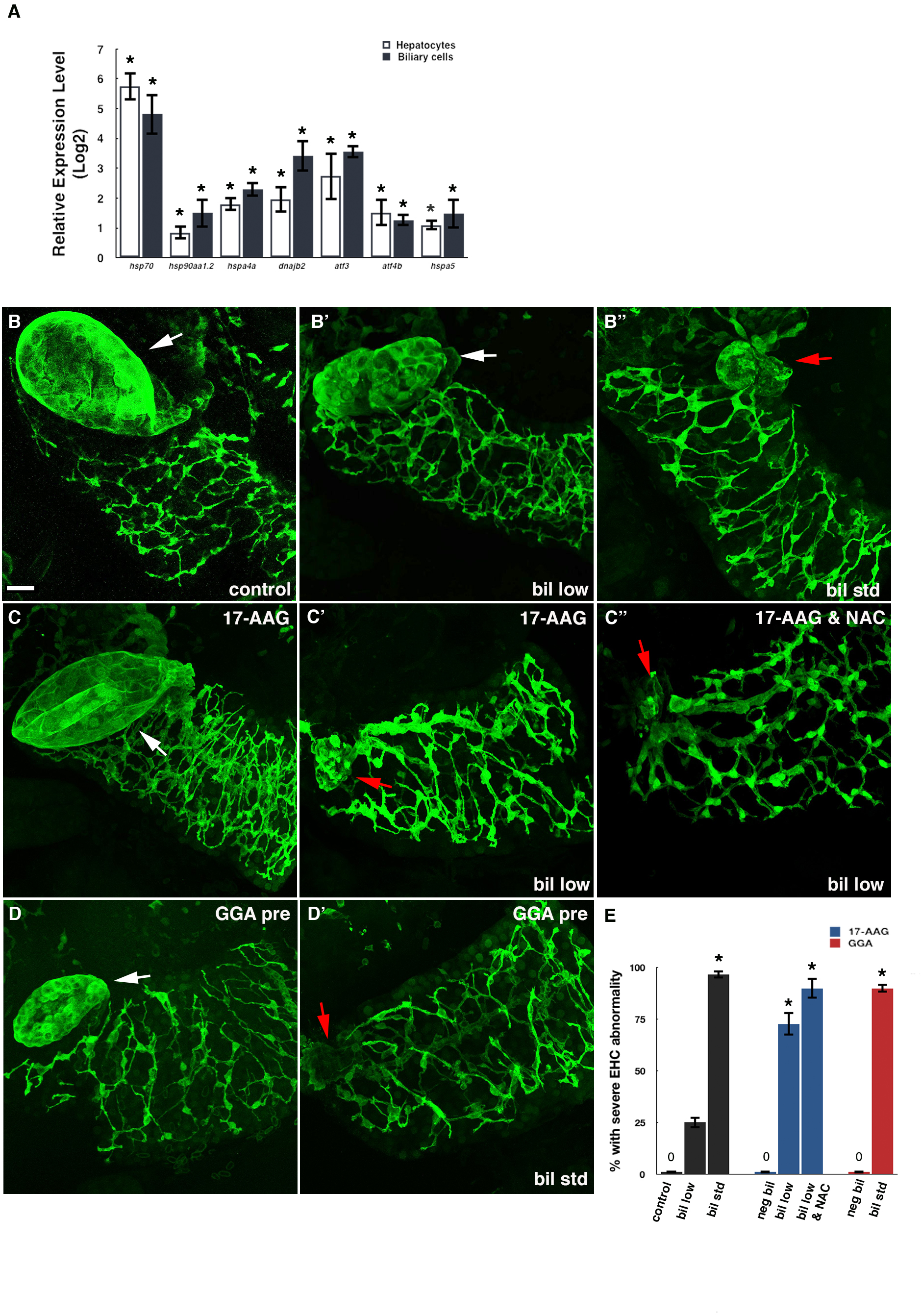
Hsp90 inhibition sensitizes EHC to biliatresone. (A) Expression of heat shock response and unfolded protein response genes in biliatresone-treated WT larval hepatocytes and cholangiocytes *versus* controls. means +/- SEM, *p<0.02 in comparison to control larvae. (B-D’) Confocal projections through the livers of larvae immunostained with anti-Annexin A4 antibody show that Hsp90 inhibition *via* 17-AAG sensitizes EHC to low dose biliatresone at 24 hr (B, B’, C, C’) that is not mitigated by NAC (C’’). Pre-treatment with the Hsp70 activator GGA fails to attenuate EHC injury induced by standard dose biliatresone. (B’’, D-D’). White arrow – gallbladder; red arrow – gallbladder remnant. Scale bar, 20 μm. (E) Percentage of WT larvae with severe EHC abnormalities after indicated exposures. *n* = 30-32 per condition, mean +/- SEM, *p<0.01 in comparison to control larvae.

### Validation of genetic variants linked to HSP90 as candidate BA risk factors

Whole exome sequencing recently identified potentially deleterious heterozygous variants in genes encoding HSP90 interactors STIP1 and REV1 in two individuals with isolated BA.^11^ STIP1 is an evolutionarily conserved co-chaperone of HSP90 and HSP70.^22^ REV1 is HSP90 client protein and functions as a Y-family polymerase that plays a central role in translesion DNA synthesis during replication stress.^23, 24^ To test the hypothesis that comparable heterozygous variants in zebrafish *stip1* or *rev1* could exacerbate biliatresone-mediated injury, we first used CRISPR-Cas9 to engineer loss-of-function mutations in both genes (Fig. S2A). Interestingly, whereas STIP1 knockout mice do not survive past E10,^25^ homozygous zebrafish *stip1* mutant larvae develop normally, but only a small percentage survive as adults (n=2 genotyped fish derived from heterozygous mating *versus* 9 predicted). Hepatobiliary development was normal in both heterozygous and homozygous *stip1* mutant larvae, however each was sensitized to low dose biliatresone (Fig. 4A-B’’, D), which was not attenuated by co-treatment with NAC (4B’’’, D). Complementary observation of impaired stress tolerance was made in the *rev1* mutants, which developed normally and survived as fertile adults (Fig. 4C-D).

**Fig. 4.**
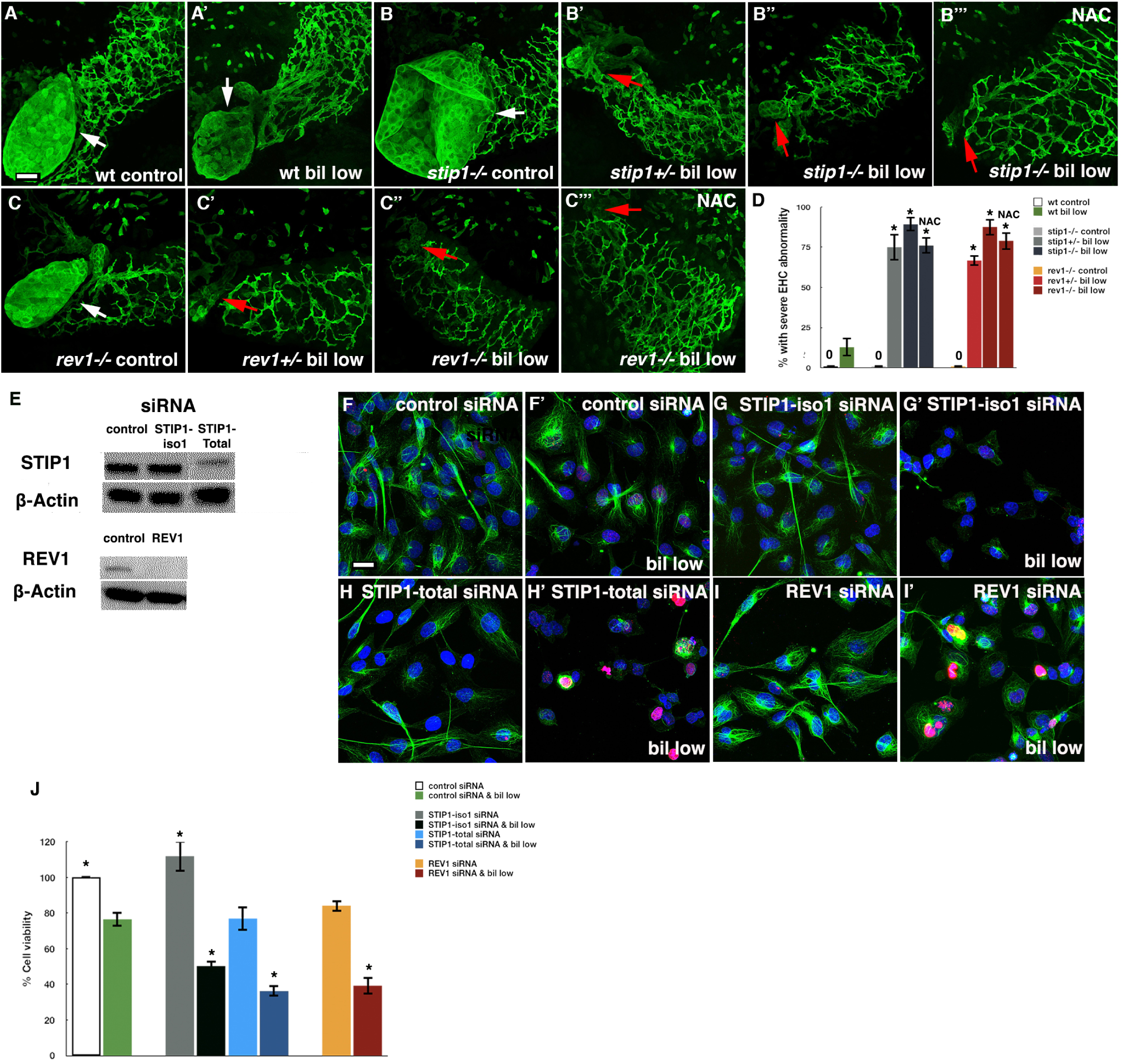
Stip1 and Rev1 disruption sensitizes zebrafish and human cholangiocytes to biliatresone. (A-C’’’) 5 dpf WT (A-A’), *stip1* (B-B’’’), *rev1* (C-C’’’) mutant larvae were exposed to DMSO *versus* low-dose biliatresone with or without NAC for 24 hr. Immunostaining with anti-Annexin A4 antibody followed by confocal microscopy show severe EHC injury in *stip1* and *rev1* mutant larvae that is not attenuated by NAC. White arrow – gallbladder; red arrow gallbladder remnant. (D) Percentage of wt, *stip1* and *rev1* larvae with severe EHC abnormalities post exposure to low-dose biliatresone with or without NAC. *n* = 30 per genotype and condition, mean +/- SEM, *p<0.01 in comparison to control larvae. (E) Western blot showing protein levels of STIP1 and REV1 extracted from H69 cells 48 hr post transfection with the indicated siRNAs. (F-I’) Transfection with siRNA targeting *STIP1-iso1* (G-G’), *STIP1*-total mRNA (H-H’) or *REV1* (I-I’), sensitizes H69 cells to low dose biliatresone, as revealed by microtubule instability (anti-acetylated tubulin antibody; green) and/or accumulation of γ-H2AX foci (red) with DAPI (blue) counterstain. Scale bar, 10 μm. (J) Relative cell viability of H69 cells exposed to low dose biliatresone at the indicated transfections as determined by MTS assay. Results expressed as a percentage of control ± SEM, *p<0.01 in comparison to H69 cells treated with control siRNA and biliatresone.

The BA-associated *STIP1* variant is a stop-gain mutation that targets a primate-specific isoform that is also the longest transcript arising from this gene (isoform 1). To get a better understanding of this isoform, we examined its expression in biliatresone-treated H69 cells. Interestingly, we found that while isoform-1 is not the predominant isoform in this cell type, its mRNA was up-regulated by 3.5 fold after 6 hr of biliatresone exposure, whereas total *STIP1* mRNA was only increased 1.5 fold. To determine if isoform-1 affects cholangiocyte responses to biliatresone, we asked whether its selective knockdown, *via* siRNA, sensitized H69 cells to a low dose of the toxin (0.5 μg/ml). RT-PCR showed significant reduction in *STIP1-iso1* mRNA levels (70.6%) with no significant effect on total *STIP1* expression (Fig. S2B). This specific knockdown caused pronounced destabilization of microtubules in biliatresone-treated cells, as previously shown in mouse cholangiocytes,^26^ along with decreased cell survival, whereas neither occurred with the control siRNA (Fig. 4F-G’, J). Similar sensitization to biliatresone was observed with siRNA that targets all *STIP1* isoforms, which led to total STIP1 reduction at both mRNA (79.5%) and protein levels (49.9%) (Fig. S2B; 4E, H, H’, J).

Loss of Stip1 triggers DNA double strand breaks in mouse embryonic fibroblast and fly ovary,^25, 27^ thus we assessed for formation of nuclear foci of phosphorylated H2AX (γ-H2AX), a sensitive indicator of DNA damage and replication stress. Interestingly, knockdown of total STIP1 protein led to induction of γ-H2AX foci in H69 cells treated with low-dose biliatresone, whereas no change was observed with isoform-1 specific STIP1 knockdown (Fig. 4G-H’). Collectively, these data suggest that while STIP1 isoform-1 may play an important role in preventing cytoskeletal collapse mediated by exogenous electrophiles such as biliatresone, the other isoforms are essential for combatting oxidant-induced DNA damage.

Similar experiments were conducted in human cholangiocytes to examine the effect of REV1 depletion on biliatresone sensitivity. Treatment of H69 with REV1-specific siRNA resulted in significant REV1 knockdown, at both the mRNA and protein levels (Fig. S2B; 4E). While unstressed REV1-deficient cells were normal, pronounced accumulation of γ-H2AX and reduced cell viability were observed in cells exposed to low-dose biliatresone (Fig. 4I-J). Of note, REV1 knockdown did not lead to any apparent microtubule instability in cells treated with low-dose biliatresone, thus demonstrating a different injury mechanism compared with STIP1 knockdown.

### Activation of cGMP signaling rescues biliatresone-mediated cholangiocyte injury in zebrafish and human cholangiocytes

The discovery that biliatresone affected PQC independently of redox homeostasis indicated that it might be possible to enhance the protective effect of NAC by activating this or other stress response pathways. To test this idea, we screened a library of known drugs (JHCC; Version 1.0) for their ability to blunt EHC damage in WT biliatresone-treated larvae. Of 1514 drugs screened, a consistent effect was only seen with two drugs, the phosphodiesterase inhibitors (PDEi) aminophylline and trequinsin. Aminophylline is a non-specific PDEi that blocks the hydrolysis of cAMP and cGMP in a broad range of cell types.^28^ Trequinsin preferentially targets PDE isoform 3 (PDE3), also leading to increased levels of cAMP and cGMP.^29^ To confirm the specificity of these results, we co-treated WT larvae with standard dose biliatresone and independently sourced PDEi for 72 hr (Fig. 5A, Table S3). These experiments confirmed the activity of aminophylline (Fig. 5C). However, the response to trequinsin was inconsistent (data not shown). Interestingly, two PDE5 inhibitors, vardenafil and tadalafil, which block cGMP degradation and are clinically approved for erectile dysfunction and pulmonary hypertension,^30^ had a comparable effect as aminophylline (Fig. 5C’, E; S3B, D).

**Fig. 5.**
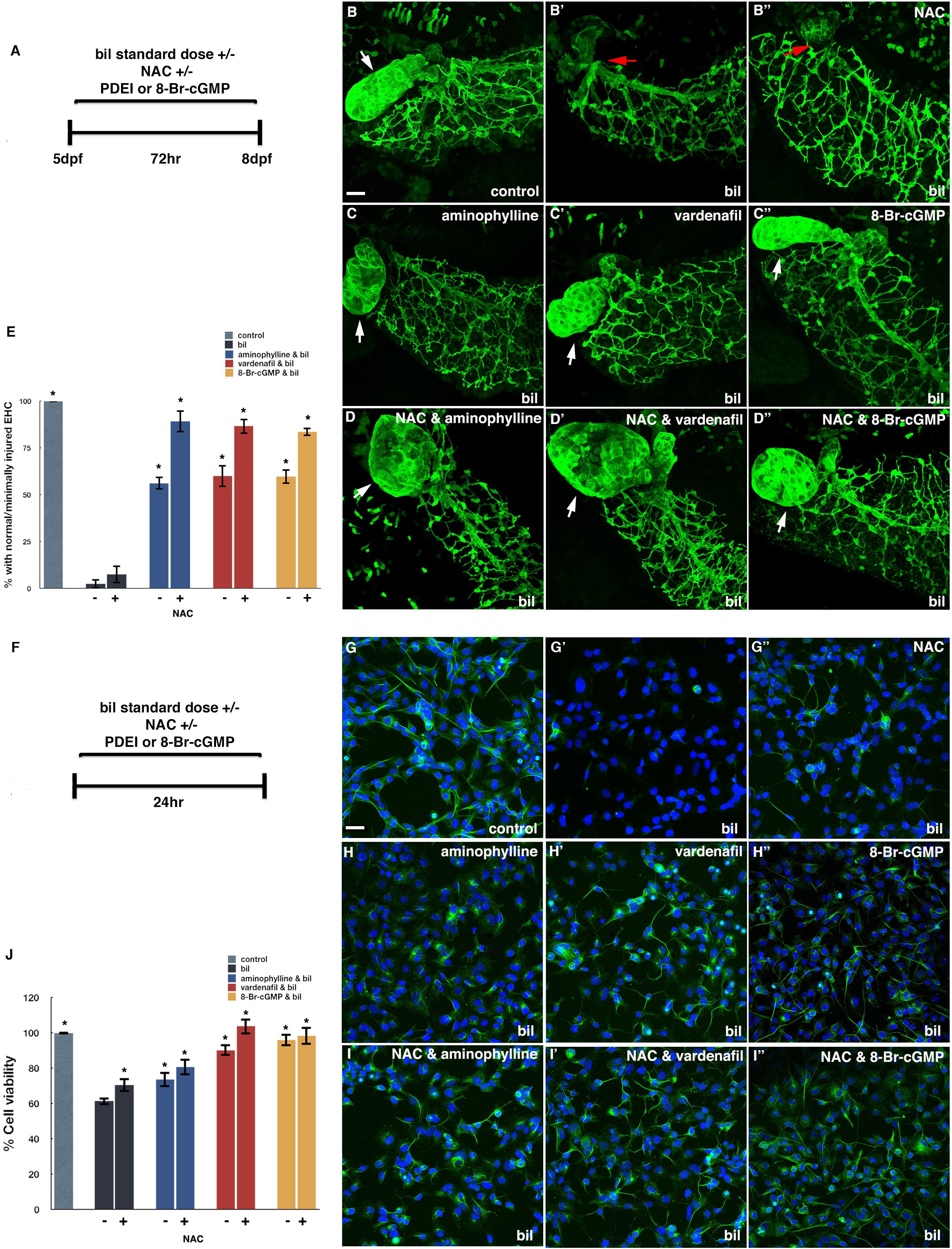
cGMP signaling attenuates biliatresone toxicity in the zebrafish and human cholangiocytes. (A) Experimental schematic showing 5 dpf WT larvae exposed to standard dose biliatresone with or without cGMP signaling enhancers (aminophylline, vardenafil, 8-Br-cGMP) +/- NAC for 72 hr. (B-D’’) Immunostaining with anti-Annexin A4 antibody followed by confocal microscopy show reduced EHC injury in biliatresone-treated larvae co-exposed to cGMP activators (C-C’’) that is further enhanced by NAC (D-D’’). NAC monotherapy not effective (B’’) at this time point. White arrow – Gallbladder; red arrow – gallbladder remnant. Scale bar, 20 μm. (E) Percentage of biliatresone-treated larvae rescued by cGMP modulators +/- NAC. *n* = 30-36 per condition, mean +/- SEM, *p<0.01 in comparison to biliatresone-treated larvae. (F) Experimental schematic showing H69 cells exposed to standard dose biliatresone with or without cGMP modulators +/- NAC for 24 hr. (G-I’’) Tubulin immunostaining (green) and nDAPI (blue) counterstain of H69 cells show cGMP enhancers with or without NAC prevent biliatresone-induced microtubule depolymerization. Scale bar, 10 μm. (H) cGMP enhancers +/- NAC improve viability of biliatresone-treated H69 cells (MTS assay). Results are expressed as a percentage of control ± SEM, *p<0.01 in comparison to biliatresone-treated H69 cells.

To confirm that PDEi-mediated rescue arose from activation of cGMP signaling, we examined the effect of the cGMP analog 8-Bromoguanosine 3’,5’-cyclic monophosphate (8-Br-cGMP) and the heme-dependent and independent activators of soluble guanylate cyclase (sGC) (BAY 41-2272 and BAY 58-2667)^31^ in biliatresone-treated larvae. All three treatments imparted resistance to biliatresone that was comparable to treatments with the PDEi (Fig. 5C’’, E; S3B’, B’’, D). In contrast, co-treatment with the adenylate cyclase activator, forskolin,^32^ which increases levels of cAMP, but not cGMP, did not reduce biliatresone-toxicity (Fig. S4).

Activation of cGMP signaling had a comparable effect on biliatresone toxicity in two human cholangiocyte cell lines (H69 and NHC) (Fig. 5F; S5). On its own, standard dose biliatresone (1.0 μg/ml) disrupted microtubules and decreased cell survival in both cell lines (Fig. 5G, G’, J; S5A, A’, D) without affecting cGMP levels (Fig. S6). Both toxicity readouts were significantly reduced by treatment with PDEi and 8-Br-cGMP (Fig. 5H-H’’, J; S5B-B’’, D). These findings confirm that activation of cGMP signaling can rescue biliatresone-mediated injury in both zebrafish and human cholangiocytes. Furthermore, they suggest the rescue does not simply restore a global deficit in cGMP.

### cGMP signaling functions independently of glutathione anti-oxidant responses

To examine the relationship of cGMP signaling to glutathione-mediated injury responses, we asked whether PDE5 inhibitors or 8-Br-cGMP enhanced the ability of NAC to delay EHC injury in biliatresone-treated zebrafish larvae. We found that activation of cGMP signaling in zebrafish significantly improved the effect of NAC, with 80-90% of larvae showing normal EHC or minimal EHC injury after 72 hr of treatment with standard dose biliatresone (Fig. 5D-E; S3C-D). NAC monotherapy, on the other hand, was not effective at this treatment time point (Fig. 5B’’, E; S3A’’, D). Comparable effects were also observed in co-treatment of the human cholangiocyte cell lines exposed to biliatresone with NAC alone improving cell viability but not preventing microtubule destabilization (Fig. 5G’’, I-J; S5A’’, C-D).

As further proof that cGMP signaling functioned independently of glutathione-mediated responses, we examined the effect of vardenafil and 8-Br-cGMP in biliatresone-treated *gclm* mutants and observed significant mitigation of both EHC and IHD injury in the mutant larvae (Fig. S7A-E). Importantly, they did so without altering total hepatic GSH levels, which were reduced comparably in the mutants regardless of whether they received vardenafil or 8-Br-cGMP (Fig. S7F).

### Inhibition of biliatresone-induced biliary injury by cGMP requires intact HSP70-HSP90 chaperone machinery

cGMP signaling has been shown to enhance proteasome-mediated degradation of misfolded proteins in cardiomyocytes.^33^ We found cGMP functioned comparably in human cholangiocytes as vardenafil or Br-cGMP significantly enhanced proteasomal activity in H69 cells co-exposed to biliatresone (Fig. 6A). In addition, cGMP modulators with and without NAC reversed upregulation of *HSP70* in biliatresone-treated H69 cells as determined by RT-qPCR, further evidence of decreased protein folding burden (Fig. 6B). To address this question with greater specificity in an *in vivo* setting, we asked if 8-Br-cGMP could ameliorate biliatresone toxicity in larvae with genetically or pharmacologically altered Hsp70/Hsp90 chaperone machinery. While we found that 8-Br-cGMP reversed sensitization of *stip1* homozygote mutants to biliatresone, the treatment failed to rescue biliatresone toxicity in zebrafish larvae co-exposed to the Hsp90 inhibitor 17-AAG, thus arguing that cGMP-mediated rescue requires an intact Hsp90 in the zebrafish. (Fig. 6C-E, G).

**Fig. 6.**
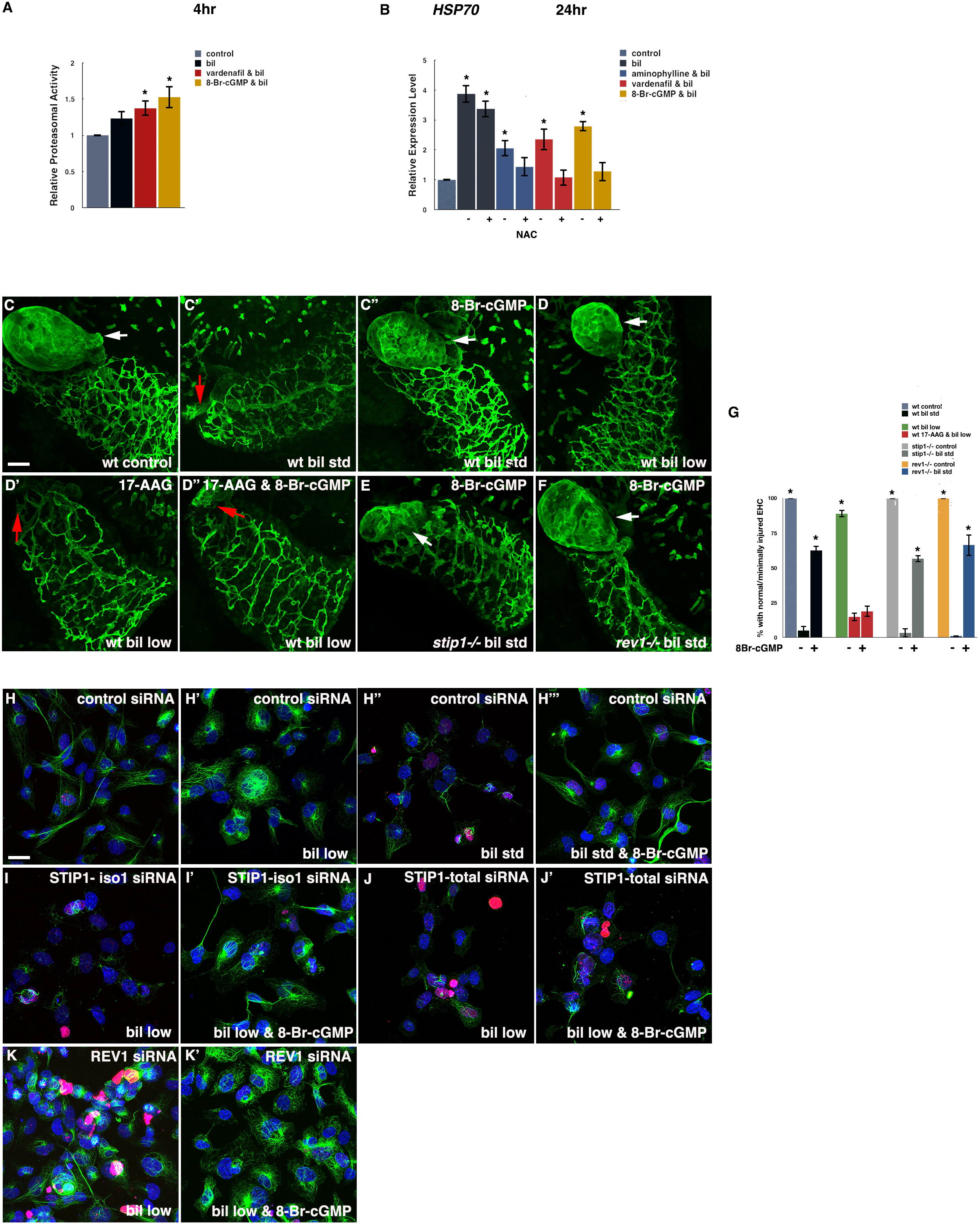
cGMP rescue of biliatresone toxicity is dependent on heat shock proteostasis. (A) Relative proteasome activity of H69 cells treated with standard dose biliatresone with or without cGMP enhancers for 4hr. (B) Relative HSP70 expression in H69 cells treated with standard dose biliatresone for 24 hr with or without cGMP enhancers and NAC. n = 3 replicates per condition, mean +/- SEM, *p<0.01 in comparison to control cells. (C-D’’) Confocal projections through livers immunostained with anti-Annexin-A4 antibody show 8-Br-cGMP fails to mitigate biliatresone toxicity in 5 dpf larvae co-treated with 17-AAG for 24 hr. (E-F) Annexin-A4 immunostaining of *stip1* and *rev1* mutants show 8-Br-cGMP attenuates biliatresone toxicity at 24 hr. White arrow – Gallbladder; red arrow – gallbladder remnant. Scale bar, 20 μm. (G) Percentage of biliatresone-treated WT, *stip1*, or *rev1* larvae with normal or minimally injured EHC following co-treatment with 8-Br-cGMP. n = 30 per genotype per condition, mean +/- SEM, *p<0.01 in comparison to larvae treated with standard-dose biliatresone. (H-K’) Tubulin immunostaining (green) with DAPI (blue) counterstain of H69 cells transfected with the indicated siRNAs followed by biliatresone with or without 8-Br-cGMP for 24 hr. Scale bar, 10 μm.

While functional Stip1 is not required for cGMP-mediated amelioration of cholangiocyte injury in the zebrafish, modulation of this co-chaperone does impact the protective effect exerted by cGMP in biliatresone-treated human cholangiocytes. Specifically, we found that 8-Br-cGMP was able to mitigate biliatresone-induced disruption of α-tubulin in cells transfected with isoform-1 specific siRNA, but not in cells with total STIP1 knockdown, arguing that cGMP in human cholangiocytes is sensitive to levels of STIP1 protein (Fig. 6H-J’).

Although cGMP activity may be dependent on an intact HSP70/90 chaperone machinery, it does not appear to rely on the HSP90 interacting protein REV1. Namely, we found that exogenous cGMP administration was able to attenuate EHD injury in *rev1* homozygous mutants and reverse γ-H2AX accumulation in biliatresone-treated REV1-depleted human cholangiocytes (Fig. 6F, G, K, K’).

### Combined use of cGMP activators and NAC blocks progressive biliary injury from biliatresone

As an environmental trigger for human BA has not been conclusively demonstrated, the timing and duration of exposure remain speculative. The recent observation that a rising total bilirubin level had high sensitivity and specificity for BA in newborn infants^34^ suggests a prenatal exposure, presumably transient, can trigger progressive and irreversible biliary damage. Supporting this model of biliary injury, we found that short exposure (6 hr) to standard dose biliatresone induced minimal or no EHC injury in zebrafish that were examined immediately after the toxin was removed, consistent with prior studies,^9^ however the injury progressed in 64.3% of larvae examined 18 hr after toxin removal (Fig. 7A-C, F). This suggests that biliatreone triggered a self-perpetuating injury cascade, and we were able to halt this injury progression by initiating combined treatment with NAC & cGMP modulators at the time of toxin removal, but not with NAC alone (Fig. 7C’-F).

**Fig. 7:**
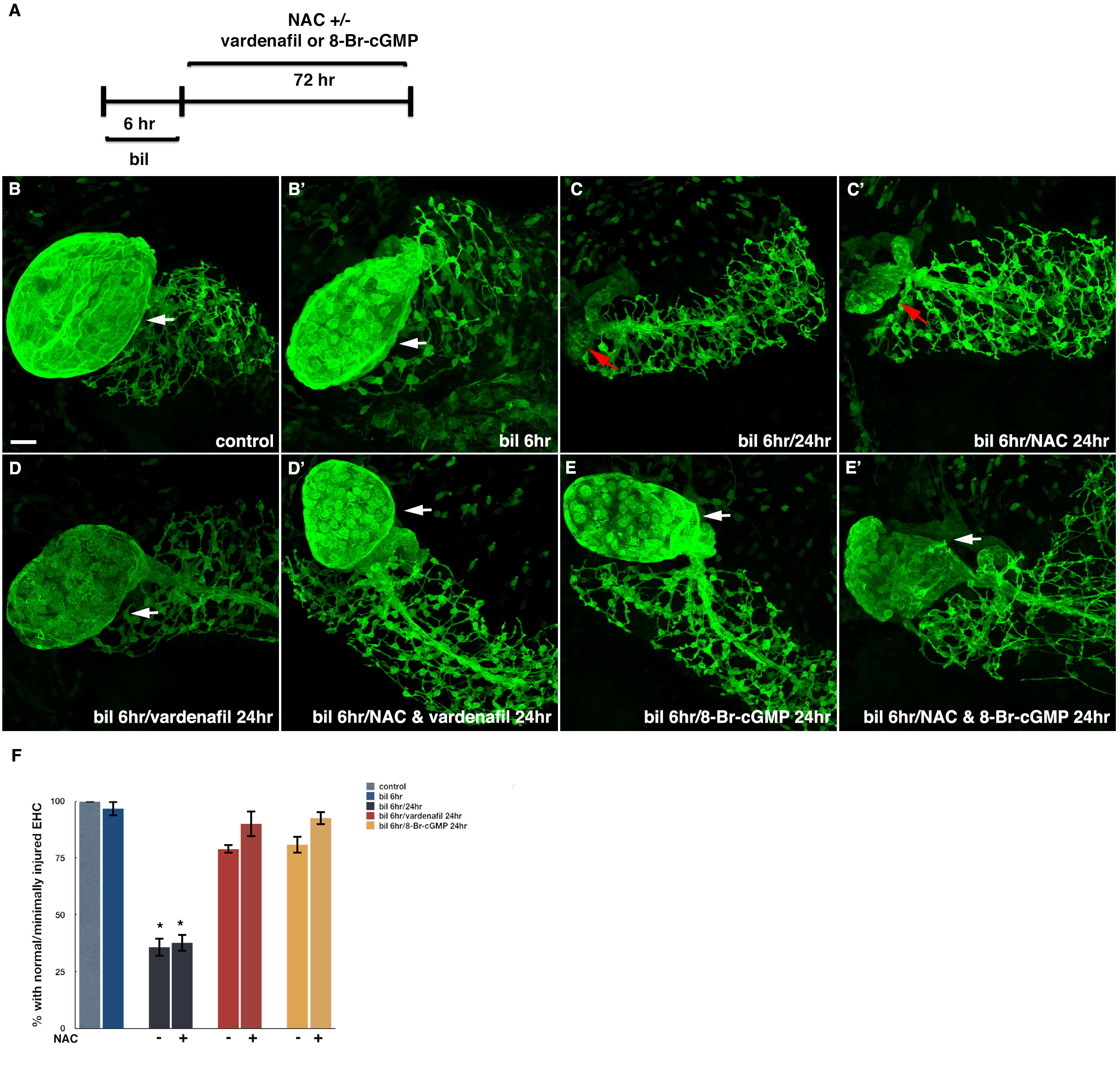
cGMP blocks progressive biliary injury from biliatresone. (A) treatment protocol with standard dose biliatresone for 6 hr, followed by cGMP modulators +/- NAC for 18 hr. (B-E’) Annexin A4 immunostaining followed by confocal microscopy showed progressive biliary injury at 18 hr after temporary exposure to biliatresone for 6hr (B-C), which was blocked by cGMP modulators +/- NAC (D-E’). (F) Percentage of larvae exhibiting normal or minimally injured EHC after biliatresone exposure for 6 hr followed by treatments with NAC/cGMP modulators. *N* = 30-35 per condition, mean +/- SEM, *p<0.01 in comparison to control larvae.

## Discussion

There is an urgent need for new treatments for BA, which despite being a rare condition, is the most common indication for pediatric liver transplantation. Unfortunately, neither the inciting disease trigger, which leads to fibrotic destruction of the extra-hepatic biliary system, nor the cause of progressive intra-hepatic bile duct disease and subsequent liver failure, has been determined.^35^ The identification of biliatresone from *Dysphania* species plants was a significant advance for the field because it was the first conclusive evidence of an environmental toxin capable of causing a biliary injury pattern that recapitulates the cardinal features of human BA.^6^

Our previous study established the importance of cholangiocyte redox heterogeneity in conferring susceptibility of EHC to biliatresone-induced oxidative injury.^9^ The work reported here provides mechanistic insights into this injury response through targeted genetic manipulation of the redox metabolism pathway coupled with *in situ* redox mapping. Specifically, we found that EHC rely on GSH synthesis and regeneration to maintain redox equilibrium under conditions of oxidative stress, as evidenced by their sensitization in *gclm, abcc2*, and *gsr* mutants. In contrast, IHC do not depend on GSH regeneration to prevent oxidative injury, but strongly depend on the re-synthesis of GSH secreted into bile by hepatocytes, as evidenced by the sensitization of *abcc2* heterozygous and homozygous mutants to biliatresone, which was reversed by the GSH precursor NAC. Conversely, NAC failed to block EHC sensitization in *gsr* mutants, indicating that despite enhanced supply of cysteine, ^16^ which is rate limiting, EHC are unable to up-regulate GSH synthesis sufficiently enough to compensate for the loss of GSH regeneration. The underlying reason for these cellular differences in maintaining redox homeostasis, whether due to differential ability to absorb GSH metabolites in bile, differential activity of synthetic enzymes, or both, is not yet known, but will be a focus of future studies.

Altogether, these data point to the possibility that gene variants affecting GSH metabolism can be otherwise silent risk factors for environmentally-induced BA in humans and affect liver recovery surgical restoration of bile flow. Supporting this, Lu et al., reported a positive correlation between transplant-free survival in BA patients and hepatic GSH metabolism gene expression measured at the time of Kasai portoenterostomy.^10^

Cellular responses to injury are complex. The protective effect imparted by the glutathione precursor NAC against biliatresone in zebrafish and mouse neonatal extrahepatic cholangiocytes reported here and in previous studies was significant but transient.^9, 26^ Transcriptional profiling data led us to examine the role of HSR in the cellular response to biliatresone as it was upregulated contemporaneously with glutathione metabolism genes. These experiments showed that disruption of HSR, either by pharmacological inhibition of HSP90, or genetically manipulating two HSP90 interacting proteins (STIP1 and REV1) linked to human BA by whole exome sequencing, sensitized zebrafish larvae and human cholangiocytes to the toxin. The sensitization of *stip1, rev1*, or larvae treated with the Hsp90 inhibitor was not reversed by NAC, and hepatic GSH level was not reduced in *stip1* mutants, thus arguing that HSR likely functions as an independent arm of the stress response to biliatresone. To our knowledge, these are the first *in vivo* studies examining epistatic interactions between redox stress and proteomic stress responses in cholangiocytes and how the two pathways might intersect in the context of biliary disease.

Previous work has linked Hsp90, Stip1 and Rev1 to mammalian stress responses in a variety of models,^25, 36^ thus we believe our observations regarding proteomic stress response elicited by biliatresone are applicable to human BA. Hepatic injury responses however are complex, and they are likely to be cell-type and context dependent. Thus, while Hsp90 inhibition exacerbated cholangiocyte injury from biliatresone and hepatic levels of HSP90 protein were significantly reduced in newly diagnosed BA infants,^37^ Hsp90 inhibition has been reported to have a protective effect in other injury models.^38^

One of the challenges in studying complex diseases lies is assigning causality to associations with genetic variants. The BA-associated *STIP1* and *REV1* variants studied here have not previously been linked to human disease, and our findings argue that heterozygous loss of function variants are normally well tolerated, however under conditions of environmental or endogenous stress, such as with exposure to a toxicant or a toxin, like biliatresone, they can predispose carriers to BA and possibly other types of biliary injury. The *STIP1* variant inactivates a primate-specific isoform that has not been studied previously, and our work here has highlighted its effect on cytoskeletal integrity upon oxidative stress exposure while the other STIP1 isoforms are needed for the DNA damage response. Collectively, these results demonstrate the crucial roles played by the HSP90 chaperone system in maintaining cytoskeletal integrity and genome stability, consistent with prior studies showing their interactions with tubulin ^39, 40^ and various DNA repair proteins.^23, 27^

The other novel finding to emerge from this study is the potential therapeutic role for cGMP signaling activators in BA. As cGMP levels were not decreased by biliatresone, the PDE5i and soluble guanylate cyclase stimulators studied here likely impart their protective effects by modulating compartmentalized cGMP signaling in cholangiocytes, which has been observed in other cell types,^41^ rather than simply restoring cGMP levels. Our studies suggest that the therapeutic benefit is mediated by the PQC machinery, at least in part by promoting proteasomal degradation of denatured proteins. Furthermore, we found that cGMP signaling depends on Hsp90, but not on the adapter protein Stip1 or the client protein Rev1, in the zebrafish larvae. This is consistent with prior studies showing the crucial role of Hsp90 in the assembly and maintenance of 20S proteasome.^42^ Future studies will be needed to elucidate the exact proteins targeted by biliatresone and how cGMP activators affect these interactions. And while our studies point to a role for these compounds in cholangiocytes, we have not excluded a contributory effect on hepatocytes or other liver cell types.

Based on the additive ameliorative effects of NAC/cGMP in the biliatresone model, we believe that additional studies to explore the use of these approved compounds in preventing BA progression are warranted. We envision two potential points of entry for the combination NAC/cGMP therapy in the timeline of BA. First, near the onset of extra-hepatic duct obstruction, which can be determined by measurement of conjugated bilirubin at birth and at 2-3 weeks of age.^34^ This approach has a high sensitivity and specificity for detecting BA, and was recently validated in a larger multi-center study.^43^ Development of a reliable biomarker of cellular stress could potentially improve the predictive value of this diagnostic strategy, which is not optimal given the rarity of BA. The second point of entry for administrating NAC/cGMP therapy would be at the usual time of diagnosis, at 1 to 2 months of age.^34^ At this stage, there is often irreversible damage to the extra-hepatic ductal system and varying degrees of portal fibrosis. The cause of progressive biliary fibrosis, even after restoration of bile flow, is not entirely know, but presumably reflects persistent activation of cholangiocyte damage pathways triggered by the inciting injury, combined with bile acid-mediated injury and secondary activation of adaptive immunity.^44, 45^ The potential therapeutic benefit of instituting NAC/cGMP combination therapy at this stage is supported by our observation using the zebrafish larvae that activation of damage pathways following a transient environmental exposure (i.e biliatresone) can be self-sustaining, but the progressive injury can be halted by employing this therapeutic strategy.

Altogether, findings from this study demonstrate that while direct consumption of biliatresone by humans is unlikely, the toxin-induced BA model can provide novel insights into cholangiocyte injury mechanisms, and through this, insights into genetic risk factors and new treatment strategies. They also point to the power of using experimental models to identify pathogenic pathways, which can then be modulated to interrogate candidate genes, hence functionally connecting genetic variations with disease-relevant phenotypes and complementing ideas that have emerged from human genetic studies and other experimental BA models.

## Abbreviations

17-AAG: 7-*N*-allylamino-17-demethoxygeldanamycin;
BA: biliary atresia;
bil: biliatresone;
8-Br-cGMP: 8-Bromoguanosine 3’,5’-cyclic monophosphate;
EHC: extrahepatic cholangiocytes;
Gcl: glutamate cysteine ligase;
GGA: geranylgeranyl acetate;
gsr: *glutathione reductase*;
GSH: reduced glutathione;
GSSG: oxidized glutathione;
IHC: intrahepatic cholangiocytes;
NAC: *N-*acetylcysteine;
PDE5i: Phosphodiesterase-5 inhibitors;
PQC: protein quality control;
WT: wild-type

