## supplementary information for "Impaired redox and protein homeostasis as risk factors and therapeutic targets in toxin-induced biliary atresia"

### Supplementary Methods

***Generation of targeted mutations in zebrafish using CRISPR/Cas9 system.*** After one-cell stage microinjection, genomic DNAs was extracted from pools of 20 one-day old embryos, and the targeted areas were amplified and sequenced using primers listed in Table S1 to confirm somatic mutations. Injected embryos were raised to adulthood (F0) and screened for the germline-transmitting founders. The F0 founders were subsequently outcrossed with WT fish to obtain the heterozygous F1 and F2 offspring.

***Drug treatments.***

Zebrafish larvae (5 dpf) reared in 24-well plates were exposed to lyophilized biliatresone resuspended in anhydrous dimethyl sulfoxide (DMSO, 0.5% in embryo medium) for 24-72 hr. The biliatresone was synthesized as previously described.[15] The larvae were either treated with low-dose biliatresone (0.25 μg/ml) or the standard dose (0.5 μg/ml) depending on the experimental condition. Additional compounds were used for the zebrafish studies at the indicated dose: *N*-acetylcysteine (NAC; Sigma-Aldrich; 20μM), aminophylline (Selleckchem, 100μM), vardenafil (Selleckchem, 1.5μM), tadalafil (Selleckchem, 1.5μM), 8-Bromoguanosine 3',5'-cyclic monophosphate (8-Br-cGMP, Sigma-Aldrich, 25μM), BAY 41-2272 (Sigma-Aldrich, 50μM), BAY 58-2667 (Sigma-Aldrich, 10μM), forskolin (Sigma-Aldrich, 0.5μM); 17-*N*-allylamino-17-demethoxygeldanamycin (17-AAG, Sigma-Aldrich, 13.2μM); geranylgeranyl acetate (GGA, Sigma-Aldrich, 20μM). The medium containing each compound was changed daily. All assays were repeated three times with at least 30 treated larvae and an equal number of control larvae in the zebrafish experiments.

Cultured cholangiocytes were serum starved for 24 hr prior to exposure to either low-dose biliatresone (0.50 μg/ml) or the standard dose (1.0 μg/ml) for 24 hr depending on the experimental condition. Additional compounds were used for the cell studies at the indicated dose: NAC (10μM), aminophylline (60μM), vardenafil (5μM), tadalafil (5μM), 8-Br-cGMP (500μM).

***Immunofluorescence.*** After chemical treatments, larvae or cells were immunostained with mouse monoclonal antibody Annexin A4 (1:100) (Abcam, Cambridge, MA), or acetylated α-tubulin (1:300) (Sigma-Aldrich), respectively, and processed for confocal microscopy as previously described.[9] Increased gain was applied to samples, if necessary, to visualize hypoplastic intrahepatic ducts in zebrafish larval liver.

***Nanostring nCounter assay and data analysis.*** Actively translating mRNA was extracted from larval hepatocytes and cholangiocytes (*Tg(fabp10a:TRAP)*; (*Tg(krt18:TRAP)*) using the translating ribosomal affinity purification (TRAP) methodology as previously described.[9] *n* = 150-300 larvae dissected per condition. Gene expression analysis was performed with the nCounter system (NanoString Technologies) as previously described.[9] The input RNA was 100 ng per sample. The nCounter system uses molecular barcodes for direct digital detection of individual target mRNA molecules. The raw data were first normalized to the positive controls provided by the manufacturer, which accounted for variations in hybridization and purification efficiency, and then to three housekeeping genes (*bactin2*, *ef1a*, *rpl19*). The experiment was done in three biological replicates and differential gene expression was calculated relative to DMSO-treated controls.

***siRNA Transfection.*** H69 cells were transfected with neg control siRNA (ON TARGET *plus* Non-targeting siRNA, Dharmacon), STIP1-iso1 siRNAs, STIP1-total siRNAs using DharmaFECT (Dharmacon) according to the manufacturer's protocol. The siRNAs with the highest efficiency were chosen for further experiments. Cells were subsequently treated with biliatresone (0.5 μg/ml) 48 hr post transfection for an additional 24 hr prior to further analyses. A 25 nM concentration of siRNA was used for all transfections.

***RNA Extraction and qRT-PCR.*** RNA was isolated from zebrafish larval liver or cultured cholangiocytes using RNeasy Micro Kit (QIAGEN). RNA samples were quantified using a Nanodrop spectrophotometer (Thermo Fisher Scientific). Synthesis of cDNA from RNA samples was carried out using the SuperScript III First-Strand Synthesis System (Thermo Fisher Scientific). Real-time RT PCR was performed using Applied Biosystems StepOnePlus and SYBR green PCR Master Mix (Thermo Fisher Scientific) with primers listed in Table S2.

***Protein Extracts and Western Blots.*** Cells were lysed in Lysis M buffer (Roche), and lysates were sonicated and pelleted. Supernatants were denatured and 30μg of protein was separated by poly-acrylamide gel electrophoresis (Invitrogen), and then transferred to nitrocellulose membranes (Invitrogen). The following antibodies were used: STIP1 (1:1000) (Atlas Antibodies), REV1 (1:1000) (Santa Cruz, CA), $\beta$-actin (1:1000) (Santa Cruz, CA), and secondary anti-rabbit IgG (H+L)-HRP conjugate (1:5000) (Invitrogen) and anti-mouse IgG (H+L)-HRP conjugate (1:5000) (Invitrogen) antibodies. Enhanced chemiluminescence images were analyzed and quantified with ImageJ.

***Cell Viability Assay.*** Cell viability assays were performed using tetrazolium compound based CellTiter 96® AQ$_{ueous}$ One Solution Cell Proliferation (MTS) assay (Promega). Biliatresone-treated H69 were co-exposed to PDEI with and without NAC. MTS assay was then performed according to the manufacturer's instruction at 24 hr post-treatments.

***Proteasome Activity Assay.*** H69 cells treated biliatresone (0.5 μg/ml, 4hr) with or without cGMP modulators were harvested. The cell pellets were resuspended in 0.5% NP-40 and homogenized by pipetting up and down followed by centrifugation at 14000 g for 10 minutes at 4 °C. 20S proteasome activity was determined using the Proteasome Activity Assay Kit (Abcam) according to manufacturer's protocol.

**Supplementary table S1: Genotyping Primer List**

| Primer ID | Sequence |
| --- | --- |
| *abcc2* forward primer | 5'-CAGGATCTGCTTTGTTTTACCC-3' |
| *Abcc2* reverse primer | 5'-TGATAGCTGTGCGAACTTTCAT-3' |
| *gclm* forward primer | 5'-GGAGAAACACTGTACCTGAGGG-3' |
| *gclm* reverse primer | 5'-ATAACTATGAGCACCGCAAACC-3' |
| *rev1* forward primer | 5'-GCAGTTTGCAGAAGCACAATAC-3' |
| *rev1* reverse primer | 5'-AGCCTGAGTAGACTTGAGGACG-3' |
| *stip* forward primer | 5’-GTATCGCAACTGAAGGATCAAGGAAAC-3’ |
| *stip1* reverse primer | 5’-TTGCCCCAGTCTGGCTTGATC-3' |

**Table S1.** List of genotyping primers.

**Supplement table S2: RT-PCR Primer List**

| Primer ID | Sequence |
| --- | --- |
| *abcc2* forward primer | 5'-TGCTCAGAGACAAGACTCGC-3' |
| *Abcc2* reverse primer | 5'-ATCCGATCTCAGACACGACG-3' |
| *gclm* forward primer | 5'-TTGACTCTGAGCTTCCCAGTG-3' |
| *gclm* reverse primer | 5'-TGATGGATGAGCAGTCCCAC-3' |
| *gsr* forward primer | 5'-GGTACCTGCGTCAATGTTGGA-3' |
| *gsr* reverse primer | 5'-TGTGCTTTTGCTCCCTCAAATC-3' |
| *zebrafish B-actin* forward primer | 5'-CAGCCATGGATGAGGAAATC-3' |
| *zebrafish B-actin* reverse primer | 5'-TCACACCATCACCAGAGTCC-3' |
| *Hsp70* forward primer | 5'-GCGGGCTCTTTAAGGCCAA-3' |

| | |
|---|---|
| *Hsp70* reverse primer | 5'-CCCCAGCAGTGTTCACACA-3' |
| *Stip1 total* forward primer | 5'-GGCTACCAGCGCTGTATGAT-3' |
| *Stip1 total* reverse primer | 5'-TAAGTGTTCGCTGAGTGCCT-3' |
| *Stip1 iso1* forward primer | 5'-TCATTGACCCATCTCAGGCTC-3' |
| *Stip1 iso1* reverse primer | 5'-TGGTGCAGTGAATGCTCGAAG-3' |
| *human B-actin* forward primer | 5'-CTGGAACGGTGAAGGTGACA-3' |
| *human B-actin* reverse primer | 5'-AAGGGACTTCCTGTAACAATGCA-3' |

**Table S2.** List of qRT-PCR primers.

**Supplement table S3**

| PDEI | PDEI Type | Specific activity | Dose | Mitigation Effect |
|---|---|---|---|---|
| Aminophylline | Non-selective | cAMP & cGMP | 250 μM | Yes |
| Cilostazol | PDE3 | cAMP > cGMP | 8 μM | No |
| Milrinone | PDE3 | cAMP > cGMP | 80 μM | No |
| Pimobendan | PDE3 | cAMP > cGMP | 10 μM | No |
| Trequinsin | PDE3 | cAMP > cGMP | 10 μM | Inconsistent |
| Roflumilast | PDE4 | cAMP | 160 nM | No |
| Tadalafil | PDE5 | cGMP | 1.5 μM | Yes |
| Vardenafil | PDE5 | cGMP | 1.5 μM | Yes |

**Table S3.** List of independently sourced phosphodiesterase (PDE) inhibitors tested in zebrafish larvae co-treated with standard dose biliatresone.

**Fig. S1. Genetic disruption of GSH metabolism genes in zebrafish.** (A) Schematic representation of the zebrafish *gclm* and *abcc2* loci along with the CRISPR target sites, protospacer adjacent motifs, and WT and indel sequences. Nucleotide changes, confirmed by cDNA sequencing, are highlighted in red. Schematic representation of the *gsr* mutant line (sa15125) obtained from ZIRC with the single nucleotide change induced by chemical mutagenesis highlighted in red. (B and C) Lateral bright field and fluorescent images of live 5 dpf wt, *gclm*$^{-/-}$, *abcc2*$^{-/-}$, *gsr*$^{-/-}$ larvae. Larvae were soaked in the fluorescent phospholipid Bodipy-C16 for 4 hours before imaging to evaluate the hepatobiliary function. (n=20 for each genotype). Scale bars, 500 µm. (D-D''') Confocal projections through the livers of 5 dpf immunostained (anti-Annexin A4 antibody) wt, *gclm*, *abcc2*, and *gsr* homozygous mutant larvae. The gallbladders are indicated by arrow. Scale bar, 20 µM. (E) Relative expression levels of the targeted transcripts in mutant larvae compared to wt siblings as determined by quantitative RT-PCR. $n = 30$ per genotype; $*p<1x10^{-5}$ by two-tailed unpaired student *t* test. (F) Hepatic GSH levels in mutant larvae compared with wt sibling controls by mass spectrometry. $n = 15$ per genotype; $*p=1.23x10^{-6}$ by two-tailed unpaired student *t* test. (G) Relative GSH levels in gallbladders dissected from adult heterozygous and homozygous *abcc2*$^{-}$ mutant larvae compared with wt sibling controls as determined by mass spectrometry. $n = 9$ per genotype; $*p<0.02$ per one-way ANOVA with Tukey's multiple comparison test. All values are mean +/- standard errors of mean (SEM).

**Fig. S2.** (A) Schematic representation of the zebrafish *stip1 and rev1* loci along with the CRISPR target sites, protospacer adjacent motifs, and WT and indel sequences. (B) mRNA levels of *STIP1-iso1, STIP1 all variants* and *REV1* from H69 cells 24 hr post transfection with control siRNA,

STIP1-iso1 siRNA, STIP1-total siRNA, or REV1 siRNA as determined by qRT-PCR. *n* = 3 per condition, mean +/- SEM, *p<0.001 by two-tailed unpaired student *t* test.

**Fig. S3. Activation of cGMP signaling attenuates biliatresone-mediated EHC injury.** (A-C'') 5 dpf WT larvae exposed to standard dose biliatresone with or without cGMP signaling enhancers (aminophylline, vardenafil, 8-Br-cGMP) +/- NAC for 72 hr. Immunostaining with anti-Annexin A4 antibody followed by confocal microscopy show reduced EHC injury in biliatresone-treated larvae co-exposed to cGMP activators (tadalafil, BAY41-2272, BAY58-2667) (B-B'') that is further enhanced by NAC (C-C''). NAC monotherapy not effective (A'') at this time point. White arrow – Gallbladder; red arrow – gallbladder remnant. Scale bar, 20 μm. (E) Percentage of biliatresone-treated larvae rescued by the indicated cGMP modulators +/- NAC. *n* = 30-36 per condition, mean +/- SEM, *p<0.01 in comparison to biliatresone-treated larvae per one-way ANOVA with Tukey's multiple comparison test.

**Fig. S4. Activation of cAMP via forskolin fails to attenuate biliatresone-mediated EHC injury.** (A-D) 5 dpf WT larvae exposed standard dose biliatresone with or without forskolin+/- NAC for 24 hr. Confocal projections through the livers of larvae immunostained with anti-Annexin A4 antibody show the failure of forskolin +/- NAC in preventing biliatresone-induced EHC injury. White arrow – Gallbladder; red arrow – gallbladder remnant. Scale bar, 20 μm. (E) Percentage of biliatresone-treated larvae exposed to forskolin with or without NAC exhibiting normal or minimally injured EHC. *n* = 30 per condition, mean +/- SEM, *p<0.01 in comparison to control larvae per one-way ANOVA with Tukey's multiple comparison test.

**Fig. S5. Enhancing cGMP signaling reduces biliatresone toxicity in NHC cells.** NHC cells immunostained with anti-acetylated tubulin antibody (green) and counter stained by DAPI (blue). (A) Control cells; (A'-A'') cells treated with standard dose biliatresone with or without NAC for 24 hr. (B-B'') Cells treated with biliatresone and cGMP modulators (aminophylline, vardenafil, 8-Br-cGMP). (C-C'') Cells treated with biliatresone, cGMP modulators and NAC. Scale bar, 10 μm. (D) Viability of biliatresone-treated NHC cells co-exposed to cGMP modulators with or without NAC as determined by MTS assay. Results are expressed as a percentage of control (DMSO-treated cells) ± SEM, *p<0.01 in comparison to biliatresone-treated NHC cells per one-way ANOVA with Tukey's multiple comparison test.

**Fig. S6.** Comparison of cGMP levels in H69 treated with standard dose biliatresone for 24 hr *versus* control cells. bil std, standard dose biliatresone

**Fig. S7.** (A-D) 5 dpf *gclm-/-* larvae exposed to low-dose biliatresone with or without vardenafil or 8-Br-cGMP for 24 hr. Immunostaining with anti-Annexin A4 antibody followed by confocal microscopy show that cGMP modulation attenuates biliary injury induced by biliatresone. White arrow – Gallbladder; red arrow – gallbladder remnant. Scale bar, 20 μm. (E) Percentage of *gclm-/-* biliatresone-treated larvae co-exposed to vardenafil or 8-Br-cGMP with normal or minimally injured EHC. *n* = 30-32 per condition, mean +/- SEM. (F) Hepatic GSH levels in *gclm-/-* biliatresone-treated larvae co-exposed to the indicated cGMP modulators compared with mutant control larvae and their WT siblings. *p<0.01 in comparison to biliatresone-treated larvae per one-way ANOVA with Tukey's multiple comparison test.

### Supplementary Fig. 1

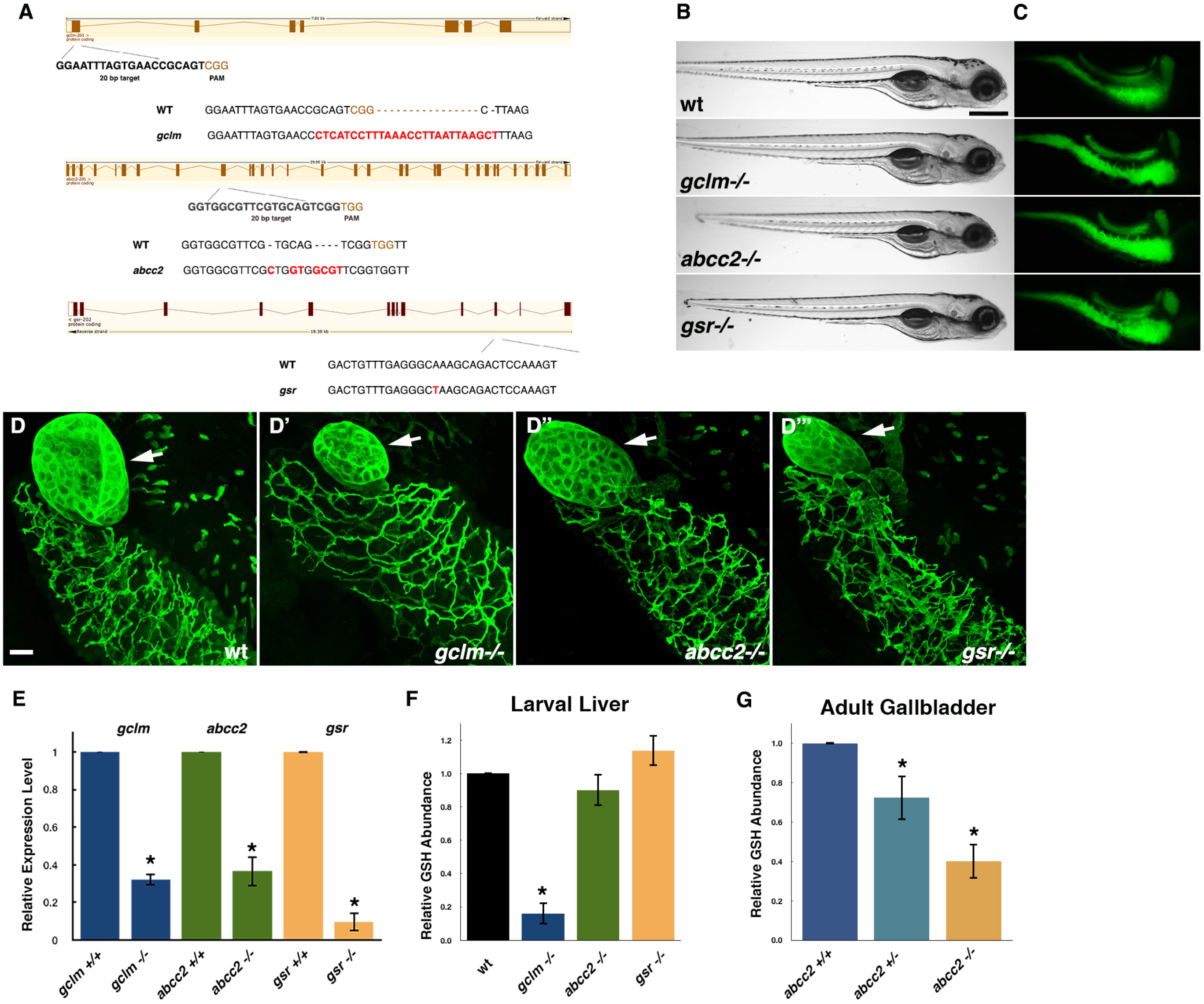

Fig. S2

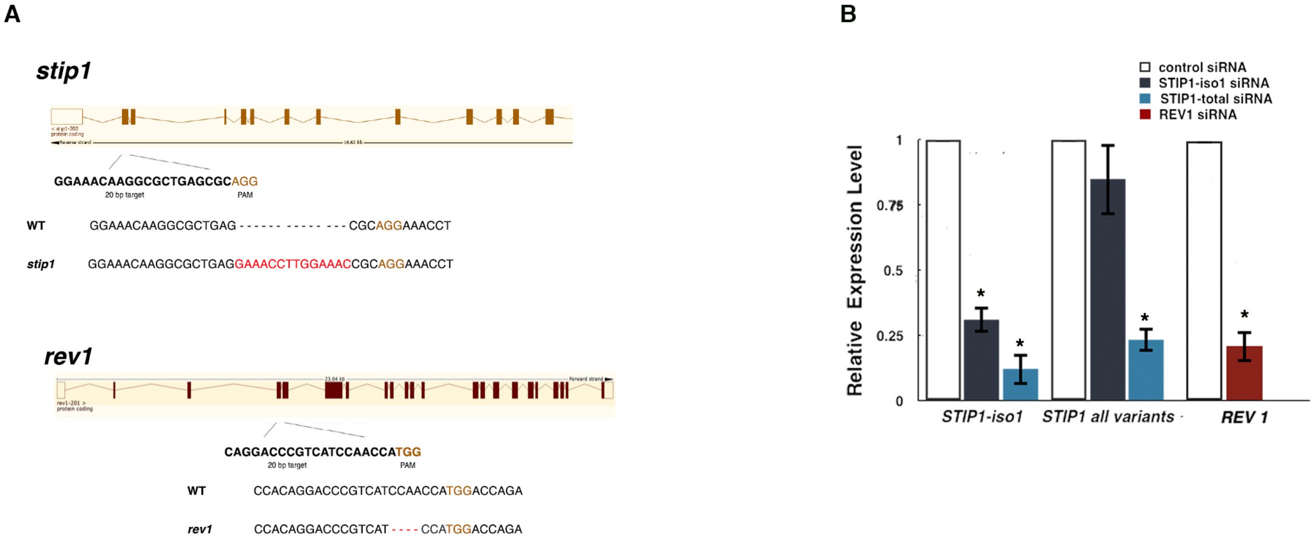
A
stip1
GGAAACAAGGCGCTGAGCGCAGG
20 bp target
PAM
WT GGAAACAAGGCGCTGAG - - - - - - - - - - - - - - CGCAGGAAACCT
stip1 GGAAACAAGGCGCTGAGGAAACCTTGGAAACCGCAGGAAACCT
rev1
CAGGACCCGTCATCCAACCATGG
20 bp target
PAM
WT CCACAGGACCCGTCATCCAACCATGGACCAGA
rev1 CCACAGGACCCGTCAT - - - - CCATGGACCAGA
B
control siRNA
STIP1-iso1 siRNA
STIP1-total siRNA
REV1 siRNA
Relative Expression Level
1
0.75
0.5
0.25
0
*
STIP1-iso1
STIP1 all variants
REV 1

**Fig. S3**

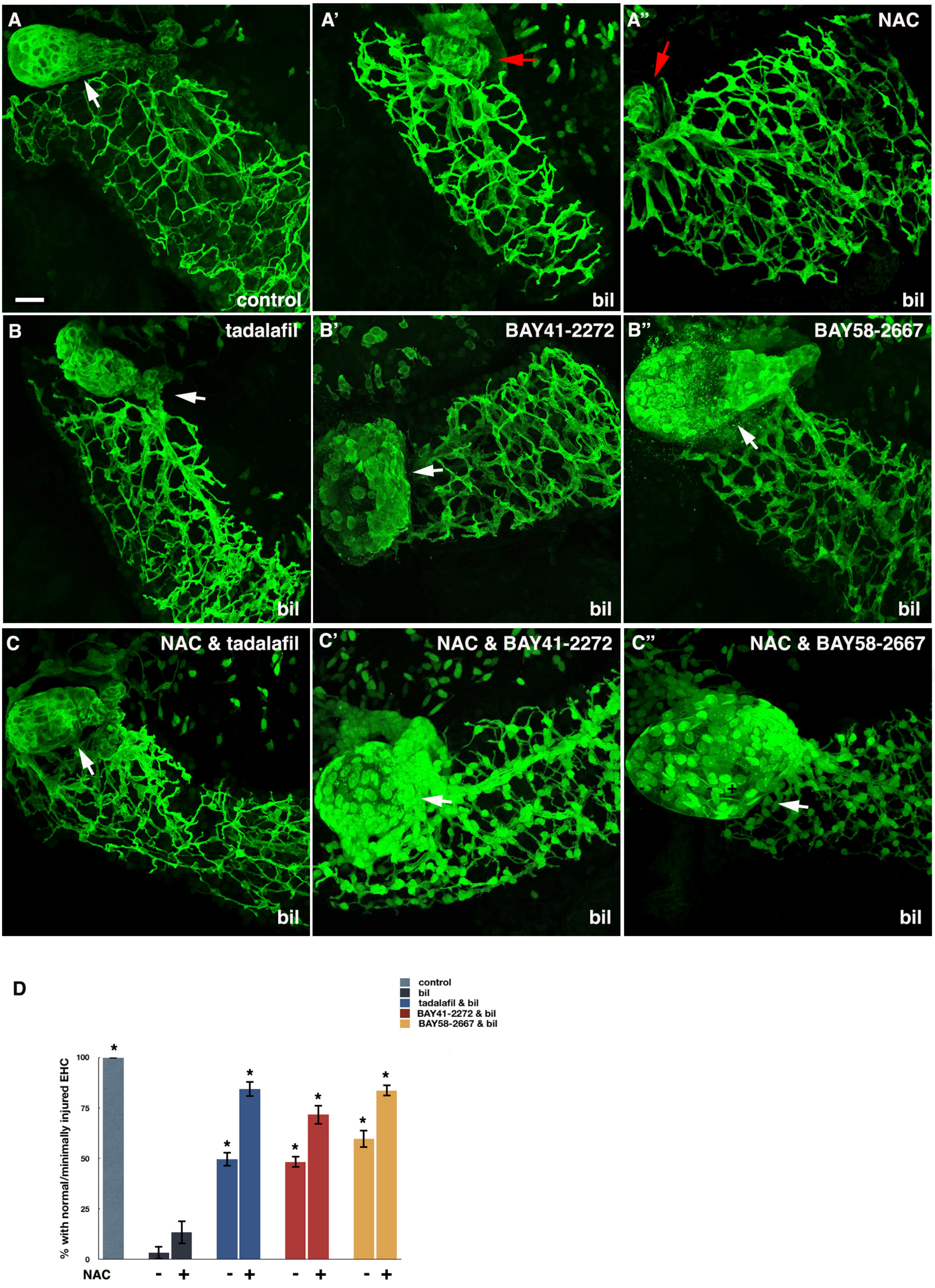
A
control
A'
bil
A''
NAC
bil
B
tadalafil
bil
B'
BAY41-2272
bil
B''
BAY58-2667
bil
C
NAC & tadalafil
bil
C'
NAC & BAY41-2272
bil
C''
NAC & BAY58-2667
bil
D
control
bil
tadalafil & bil
BAY41-2272 & bil
BAY58-2667 & bil
% with normal/minimally injured EHC
100
75
50
25
0
NAC
-
+
-
+
-
+
-
+

Fig. S4

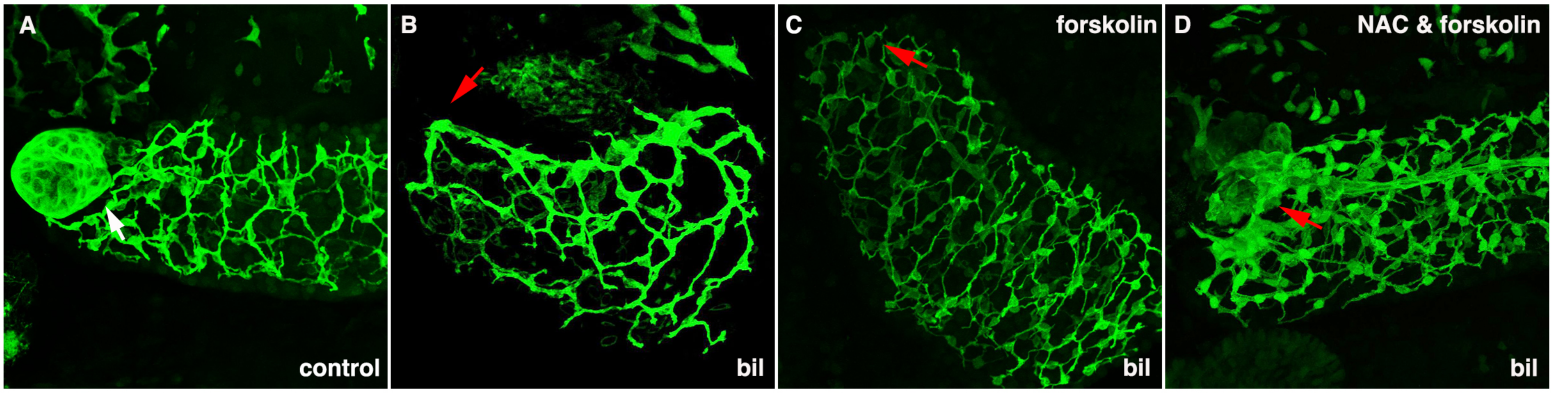
A
control
B
bil
C
forskolin
bil
D
NAC & forskolin
bil

E

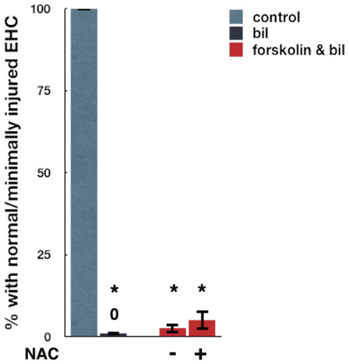
% with normal/minimally injured EHC
100
75
50
25
0
control
bil
forskolin & bil
*
0
*
*
NAC
-
+

Fig. S5

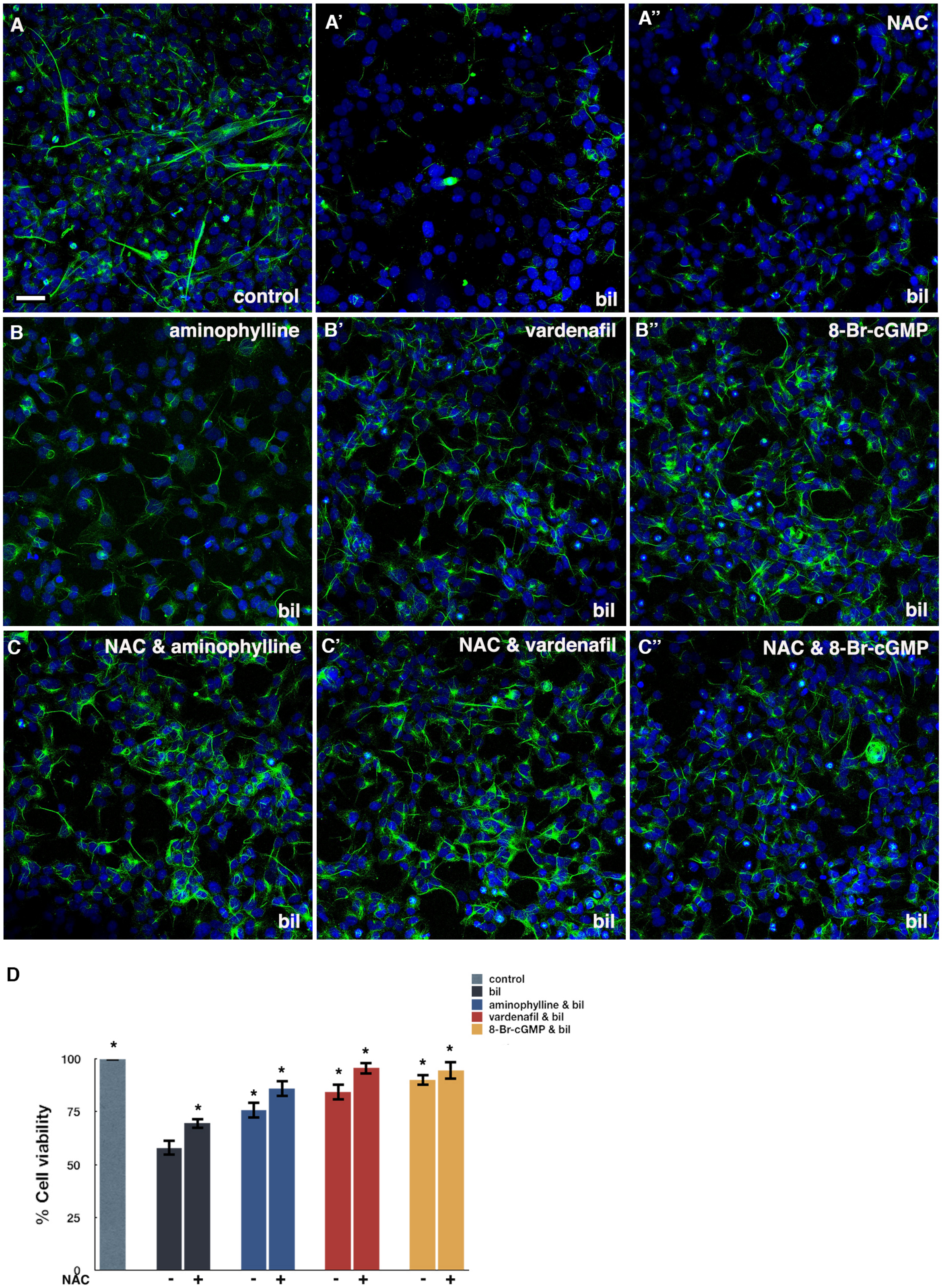
A
control
A’
bil
A”
NAC
bil
B
aminophylline
bil
B’
vardenafil
bil
B”
8-Br-cGMP
bil
C
NAC & aminophylline
bil
C’
NAC & vardenafil
bil
C”
NAC & 8-Br-cGMP
bil
D
control
bil
aminophylline & bil
vardenafil & bil
8-Br-cGMP & bil
% Cell viability
100
75
50
25
0
NAC
- + - + - + - +
*

Fig. S6

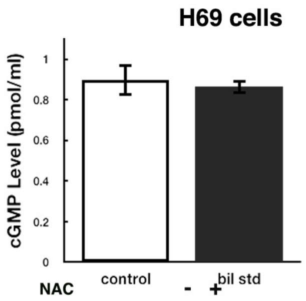

H69 cells
cGMP Level (pmol/ml)
1
0.8
0.6
0.4
0.2
0
control
bil std
NAC
-
+


Fig. S7

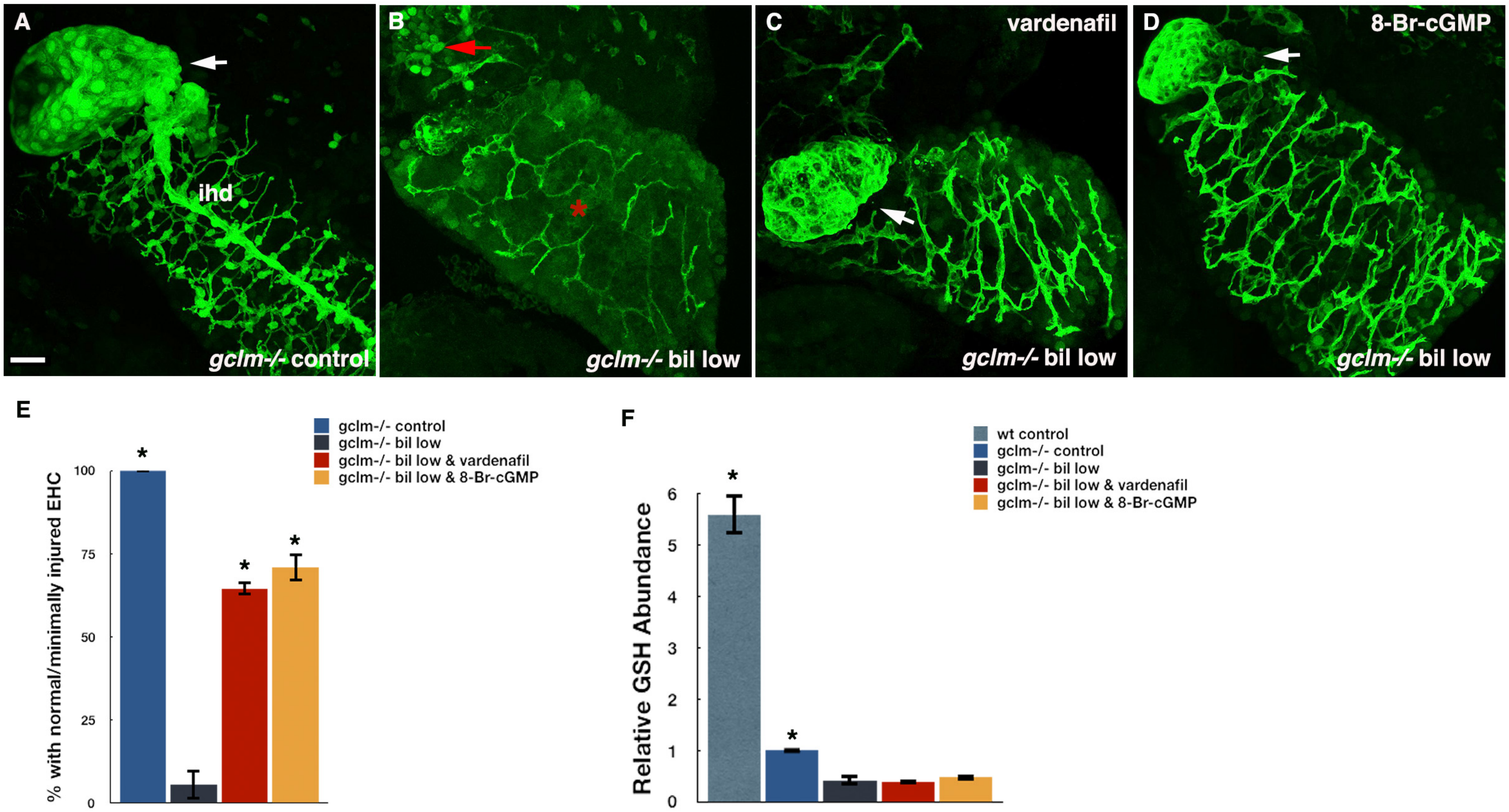

A
ihd
gclm-/- control
B
gclm-/- bil low
C
vardenafil
gclm-/- bil low
D
8-Br-cGMP
gclm-/- bil low
E
% with normal/minimally injured EHC
100
75
50
25
0
gclm-/- control
gclm-/- bil low
gclm-/- bil low & vardenafil
gclm-/- bil low & 8-Br-cGMP
F
Relative GSH Abundance
6
5
4
3
2
1
0
wt control
gclm-/- control
gclm-/- bil low
gclm-/- bil low & vardenafil
gclm-/- bil low & 8-Br-cGMP
